# Plant-associated microbiomes promote nutrient turnover in impoverished substrates of a biodiversity hotspot

**DOI:** 10.1101/2021.07.30.454538

**Authors:** Antonio P. Camargo, Rafael Soares Correa de Souza, Juliana Jose, Isabel R. Gerhardt, Ricardo A. Dante, Supratim Mukherjee, Marcel Huntemann, Nikos C. Kyrpides, Marcelo F. Carazzolle, Paulo Arruda

## Abstract

The substrates of the Brazilian *campos rupestre*s have extremely low concentrations of key nutrients, mainly phosphorus, imposing severe restrictions to plant growth. Regardless, this ecosystem harbors enormous biodiversity which raises the question of how nutrients are cycled and acquired by the biosphere. To uncover the nutrient turnover potential of plant-associated microorganisms in the *campos rupestre*s, we investigated the compositions and functions of microbiomes associated with two species of the Velloziaceae family that grow over distinct substrates (soil and rock). Amplicon, metagenomic, and metagenome-assembled genome sequence data showed that the *campos rupestres* harbor a novel assemblage of plant-associated prokaryotes and fungi. Compositional analysis revealed that the plant-associated soil and rock communities differed in taxonomic structure but shared a core of highly efficient colonizers that were strongly coupled with nutrient mobilization. Investigation of functional and abundance data revealed that the plant hosts actively recruit communities by exuding organic compounds and that the root-associated microbiomes possess a diverse repertoire of phosphorus turnover mechanisms. We also showed that the microbiomes of both plant species encompass novel populations capable of mobilizing nitrogen and that the substrate strongly influences the dynamics of this cycle. Our results show that the interplay between plants and their microbiomes shapes nutrient turnover in the *campos rupestres*. We highlight that investigation of microbial diversity is fundamental to understand plant fitness in stressful environments.

## Background

The Brazilian *campos rupestres* constitute a grassland ecoregion located on the geologically old rocky outcrops of the central and eastern regions of Brazil (Figure 1A, left). Campos *rupestres* soils are shallow, acidic, and severely nutrient-impoverished, especially low in phosphorus (P), imposing a high cost for nutrient acquisition to resident plants [1]. Nevertheless, despite this abiotic constraint, the *campos rupestres* constitute a biodiversity hotspot with an average species density among the world’s highest, harboring thousands of endemic vascular plant species from highly specialized and phylogenetically clustered lineages [2, 3].

**Figure 1.**
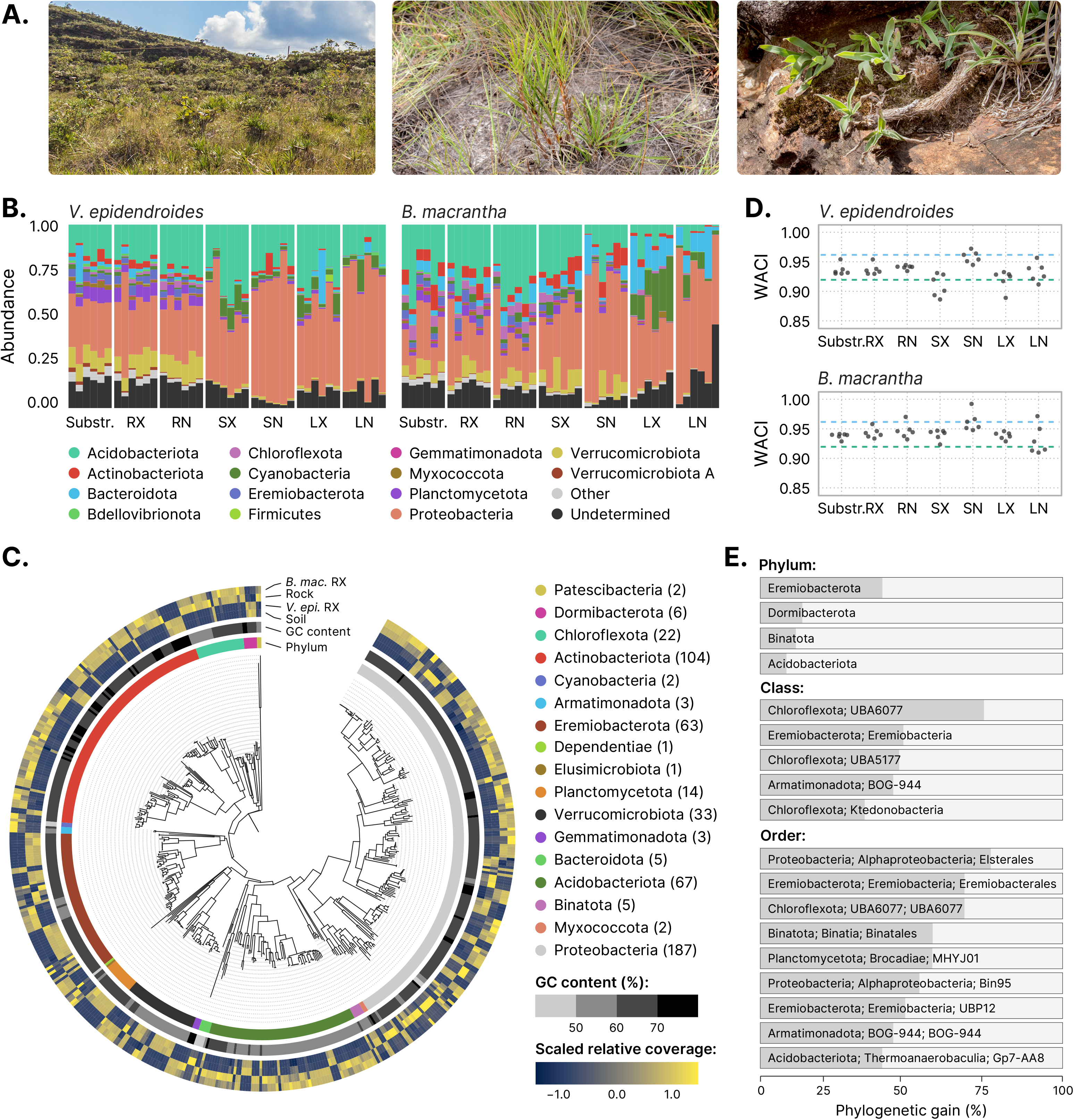
**(A)** Sampling was conducted in the *campos rupestres* grasslands ecoregion (left). *Vellozia epidendroides* (center) specimens were collected in patches of shallow soil. *Barbacenia macrantha* (right) was found in a rocky area, where it grows over exposed rocks. **(B)** Community composition inferred from 16S ASVs at the phylum level. Samples were grouped according to their environment. Bar heights are proportional to the relative abundance of the phylum. Low abundance phyla (relative abundance < 2%) were grouped under the “Other” category. **(C)** Maximum-likelihood phylogenetic tree of the bacterial MAGs presented in this study, rooted at the Patescibacteria clade. The innermost ring indicates the phylum associated with each node. The center ring shows the genomic GC content. The outermost ring represents the scaled means of log-transformed relative genomic coverages across the four environments. **(D)** Weighted average community identity (WACI) computed from 16S ASV data. The blue and green dashed lines represent the median intra-rank 16S identity at the genus and family levels, respectively. **(E)** Phylogenetic gain (PG) contributed by the MAGs to different taxa at the phylum, class, and order levels. Only taxa with PG higher than the following cut-offs are shown: 5% at the phylum level, 30% at the class level, and 40% at the order level. RX=root (external), RN = root (internal), SX = stem (external), SN=stem (internal), LX=leaf (external), LN=leaf (internal).

Plant adaptations to the nutritional scarcity of the *campos rupestres* substrates have been extensively studied. Members of the highly successful Velloziaceae family have been shown to use multiple strategies to cope with nutrient limitation, such as the formation of durable and well-defended structures [4], the efficient remobilization of P from senescent leaves, and the development of specialized radicular systems that enhance nutrient uptake via the secretion of carboxylates [5–7]. As a result of the plant-centric approach that has dominated the study of the *campos rupestres* nutritional dynamics, it remains obscure how plant growth is influenced by interactions with microorganisms.

Plant microbiomes play a fundamental role in shaping the host responses to biotic and abiotic stresses and modulating plant phenotypic plasticity. These microorganisms can form complex and stable associations that can determine plant speciation, geographic distribution patterns, and diversity to a much stronger degree than previously acknowledged [8]. This scenario implies that plants cannot be perceived as isolated entities but rather as a unit formed by the host and its associated microbiome [9]. Such tight interactions indicate that these microbial communities are not random assemblages of microorganisms, and that intricate inter-species relationships shape the microbiome composition. This molecular interplay between the plant and its microbiome has dramatically shaped the genome of root-associated bacteria [10], providing further evidence of the evolutionary interdependence of the host-microbiome system.

Microbiome-driven processes involved in nutrient acquisition are thought to have a decisive influence in naturally stressful environments, where plants are inclined to be more dependent on microbial communities for nutrient uptake than in nutrient-rich habitats, where resources are readily available [11]. In grasslands and savannahs, for example, nitrogen-fixing bacteria and mycorrhizal association contribute up to 20% of total nitrogen acquired by the vegetation [12]. In boreal forests, where nutrient availability is severely affected by low temperatures and soil pH, plant-associated microbiomes were estimated to be responsible for most of the nitrogen and phosphorus acquired by plants [13, 14].

Previous reports showing that most *campos rupestres* plants growing under P-limited substrates do not associate with mycorrhizal fungi [15] have strengthened the belief that these plants rely solely on their own adaptations to acquire phosphorus. However, a high-throughput assessment of the diversity and functions of plant-associated microorganisms has never been conducted in the *campos rupestres*, hindering any attempts to perform a comprehensive evaluation of how microbial communities influence the acquisition and turnover of nutrients in this environment.

Here, we aimed to uncover the diversity and functional roles of microbial communities associated with two species of Velloziaceae growing in two distinct nutrient-impoverished substrates: *Vellozia epidendroides* Mart. ex Schult. & Schult. f. and *Barbacenia macrantha* Lem, which are found growing in shallow soil patches (Figure 1A, center) and over rocks (Figure 1A, right), respectively. We characterized the microbiomes associated with roots, stems, and leaves of these two species, and their soil or rock substrates, and showed that novel and diverse communities associate closely with their hosts and are involved in phosphorus and nitrogen turnover and plant nutrition.

## Results

### Taxonomic and functional landscapes of Velloziaceae-associated microbiomes

Prokaryotic and fungal communities associated with *V. epidendroides* and *B. macrantha* were assessed through amplicon sequencing of the 16S gene and ITS2 region, respectively. From the substrate and plant organ samples we identified 29,008 16S and 9,153 ITS unique amplicon sequence variants (ASVs), which were assigned to three archaeal, 38 bacterial (Figure 1B), and 13 fungal (Suppl. Figure 1A) phyla. In addition to the amplicon datasets, sequence data from 16 metagenomes, comprising 25.8 Gbp, were assembled from soil-, rock-, and root-associated samples (Suppl. Table 1). From these assemblies, 522 metagenome-assembled genomes (MAGs) were recovered and assigned to one archaeal and 17 bacterial phyla (Figure 1C, Suppl. Table 2). Further clustering based on average nucleotide identity (ANI) [16] revealed 331 species-level clusters (≥ 95% ANI). The metagenomes recovered 16.3% to 42.4% of the total sequence diversity across different samples, while MAGs recovered from 10.8% to 39.1%. Additionally, most unassembled sequences were from populations closely related to those present in the metagenomes; sequence complexity was the main factor hindering metagenomic assembly (Suppl. Note 1, Suppl. Table 3).

To improve the recovery of the protein sequence space encompassed by these microbiomes, we performed protein-level assembly directly from sequence reads [17]. Clustering these peptides with the complete set of proteins predicted from the metagenomes at multiple sequence identity levels (Suppl. Table 4) revealed that the *V. epidendroides-* and *B. macrantha*-associated microbial communities harbored diverse genic repertoires with over 49 million unique proteins and six million clusters (at 50% identity).

### Taxonomic and functional novelty of *campos rupestres* microbiomes

Taxonomy could not be assigned at the family level to 46.5% of 16S and 77.5% of ITS ASVs and at the phylum level to 25.8% of 16S and 48.7% of ITS ASVs (Figure 1B, Suppl. Figure 1A). To quantify the community-level taxonomic novelty we devised the weighted average community identity (WACI) metric, which represents the abundance-weighted average identity of alignments between ASVs and their best matches in a reference database. Bacterial communities exhibited WACI around 91.8% to 95.8% (10th and 90th percentiles, Figure 1D), suggesting that they are dominated by novel genera and families (median within-rank identity of 96.4% and 92.3%, respectively [18]). There were no significant differences between the WACI of below-ground (substrate and root) and above-ground (stem and leaf) bacterial communities. However, below-ground fungal communities were significantly more novel in both plants (Suppl. Figure 1B, LMM *p*-value < 0.05; *V. epidendroides* ω^2^ ≈ 0.64; *B. macrantha* ω^2^ ≈ 0.55) and had lower WACI (median: 87.3% in *V. epidendroides* and 90.8% in *B. macrantha*).

Phylogenetic novelty was measured by computing the phylogenetic gain (PG; defined as the total branch length added by a set of genomes to a clade) brought by the MAGs to their lineages (Figure 1E). Out of the 522 genomes, 268 (51.3%) belonged to novel genera, 66 (12.6%) belonged to new families, and 19 (3.6%) were assigned to new orders. These data contributed a substantial amount of phylogenetic novelty to known taxa (Suppl. Table 5), including some understudied phyla such as the Eremiobacterota, Dormibacterota, and Binatota, that were significantly expanded (PG: 40.4%, 13.9%, and 11.7%). Surprisingly, a substantial amount of phylogenetic diversity was added to Acidobacteriota (PG: 8.6%), a phylum consistently found among abundant taxa in soils across the world [19–21]. We also found that the Elsterales, an order within the Proteobacteria that was previously found to be highly enriched in P limited soils [22], was greatly expanded (PG: 76.3%).

The functional uniqueness of the metagenomes was appraised by comparing their clustered protein repertoires to known protein families (Pfam, TIGRFAM, or KO). Around 72.3% of the non-singleton protein clusters (sequence identity ≥ 90%) contained at least one member assigned to a known family. This proportion decreased to 50.1% in clusters comprising more distantly related proteins (sequence identity ≥ 50%) (Suppl. Table 6). Additionally, functional novelty was quantified in an annotation-independent manner by comparing MAGs to closely related genomes to infer groups of orthologous genes (orthogroups) that were exclusive to the *campos rupestres* genomes at the family level. Around 563 exclusive orthogroups (comprising 4,829 genes) were found in 310 MAGs belonging to 44 families. The distribution of these orthogroups across the entire Bacteria domain revealed that 148 orthogroups (988 genes) could not be found in any other genome and likely represented lineage-specific protein families (Suppl. Note 2, Suppl. Figure 2, Suppl. Table 7).

### Comparison of the *V. epidendroides* and *B. macrantha*-associated communities

Assessment of 16S ASV data of *V. epidendroides* and *B. macrantha* microbiomes revealed that the average alpha diversity (Pielou’s equitability index [23] and richness) did not significantly differ between the prokaryotic communities associated with the two plants when discounting the effects of each tissue (Suppl. Figure 3A, B). Conversely, *V. epidendroides*-associated fungal communities exhibited significantly higher alpha diversity according to both estimates (LMM *p*-value < 0.05; richness ω^2^ ≈ 0.39; equitability ω^2^ ≈ 0.3).

Comparisons of community structure showed that the microbiomes associated with *V. epidendroides* and *B. macrantha* significantly differed (Figure 2A, Suppl. Figure 3C, D; PERMANOVA *p*-value < 0.001). Indeed, most 16S and ITS ASVs were exclusive to one or the other plant in all sample types (Figure 2B, Suppl. Figure 3E). The differences in 16S ASV abundances between the two plant-associated communities revealed that several families were significantly enriched in the microbiomes of one of the plants (Figure 2C), indicating that the differentiation was taxonomically structured. Remarkably, several Actinobacteriota families were exclusively enriched in *B. macrantha*. This structured differentiation was also verified in the MAGs, as phylogenetic proximity was significantly correlated with abundance profile similarity (Mantel test *p*-value < 0.001, Figure 1C).

**Figure 2.**
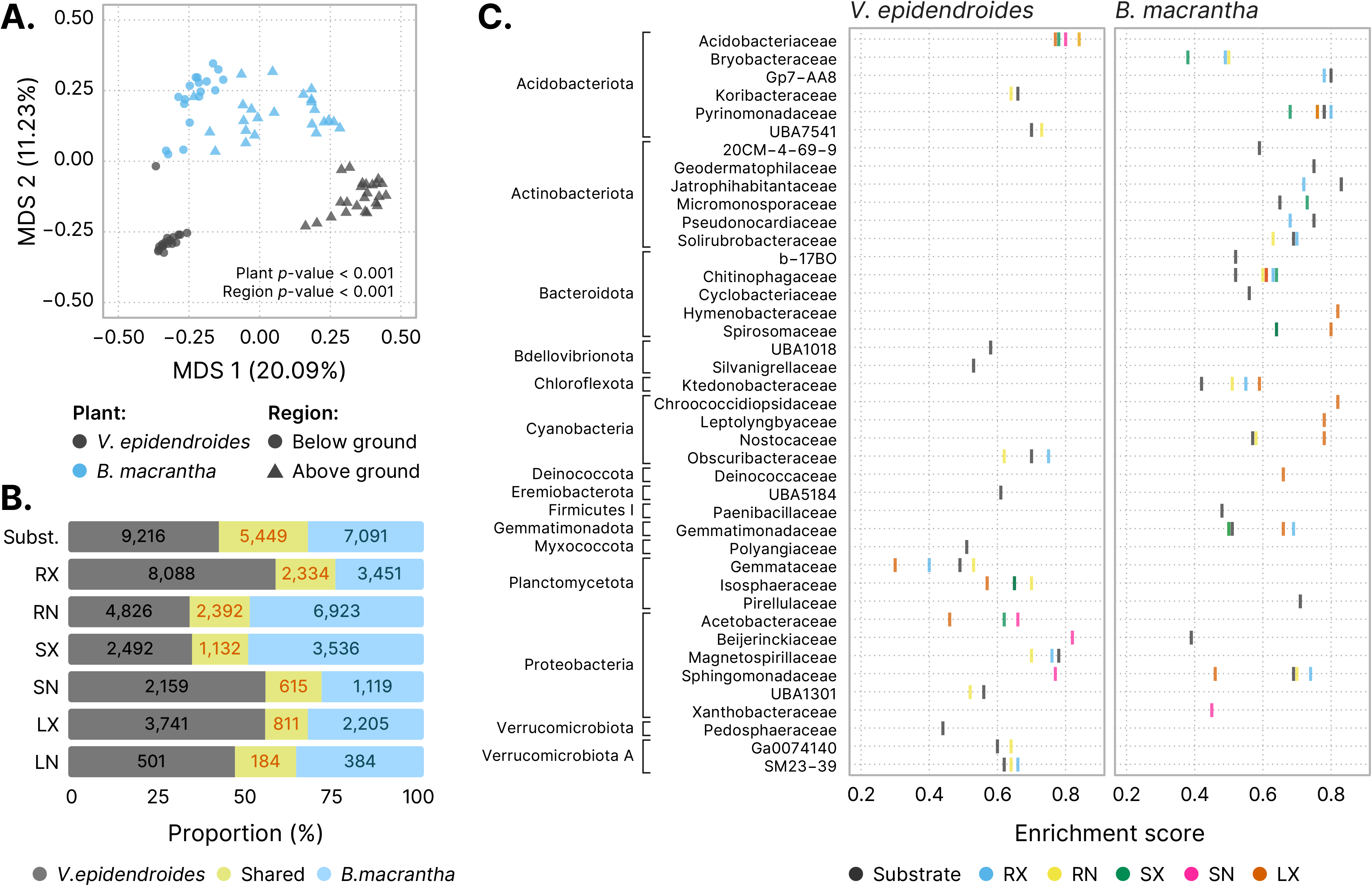
**(A)** Multidimensional scaling of the Bray-Curtis dissimilarities computed from 16S ASV abundance data. Samples are colored according to their associated plant species, and shape indicates whether they were from below ground (substrate and root) or above ground (stem and leaves) environments. The *p*-values of the groupings were obtained from PERMANOVA tests. **(B)** Bar plots representing the fraction of *V. epidendroides*-exclusive, shared, and *B. macrantha*-exclusive 16S ASVs across all sample types. The absolute numbers of ASVs within each group are shown. **(C)** Enrichment of bacterial families (grouped by phyla) in one or the other plant across all environments (circle colors). The enrichment score in the x-axis was computed using the Kolmogorov–Smirnov test and represent the deviation from a null model where ASVs from a given family are uniformly distributed in a list ranked by the ratio between the abundances in each plant. No family was found to be enriched in the internal leaf communities of either plant. RX = root (external), RN = root (internal), SX = stem (external), SN = stem (internal), LX = leaf (external), LN = leaf (internal).

Divergence in expression of the genetic repertoires of the microbiomes of the two plants was verified by assessing protein cluster abundances in each sample (Suppl. Figure 3F, G). Accordingly, 17.8% of the gene clusters had significantly different measures for average number of copies per genome (absolute log2 fold change ≥ 0.5 and s-value < 0.005) and several KEGG modules and pathways were significantly enriched in one or the other plant (116 and 95 modules, 149 and 95 pathways in *V. epidendroides* and *B. macrantha*, respectively).

Considering that both plant species cope with multiple environmental stresses, we inquired whether they shared any microbial taxa. Despite the extensive compositional differentiation, *V. epidendroides* and *B. macrantha* shared a fraction of both 16S (12% to 25%, Figure 3A) and ITS (4.2% to 25.3%, Suppl. Figure 3H) ASVs. Interestingly, the shared microorganisms tended to have higher average abundances across most tissues, often comprising more than half of the total community abundance. Additionally, several bacterial families were enriched within the shared 16S ASV sets, suggesting that some lineages had a core microbiome that could efficiently colonize both plants (Figure 3B).

**Figure 3.**
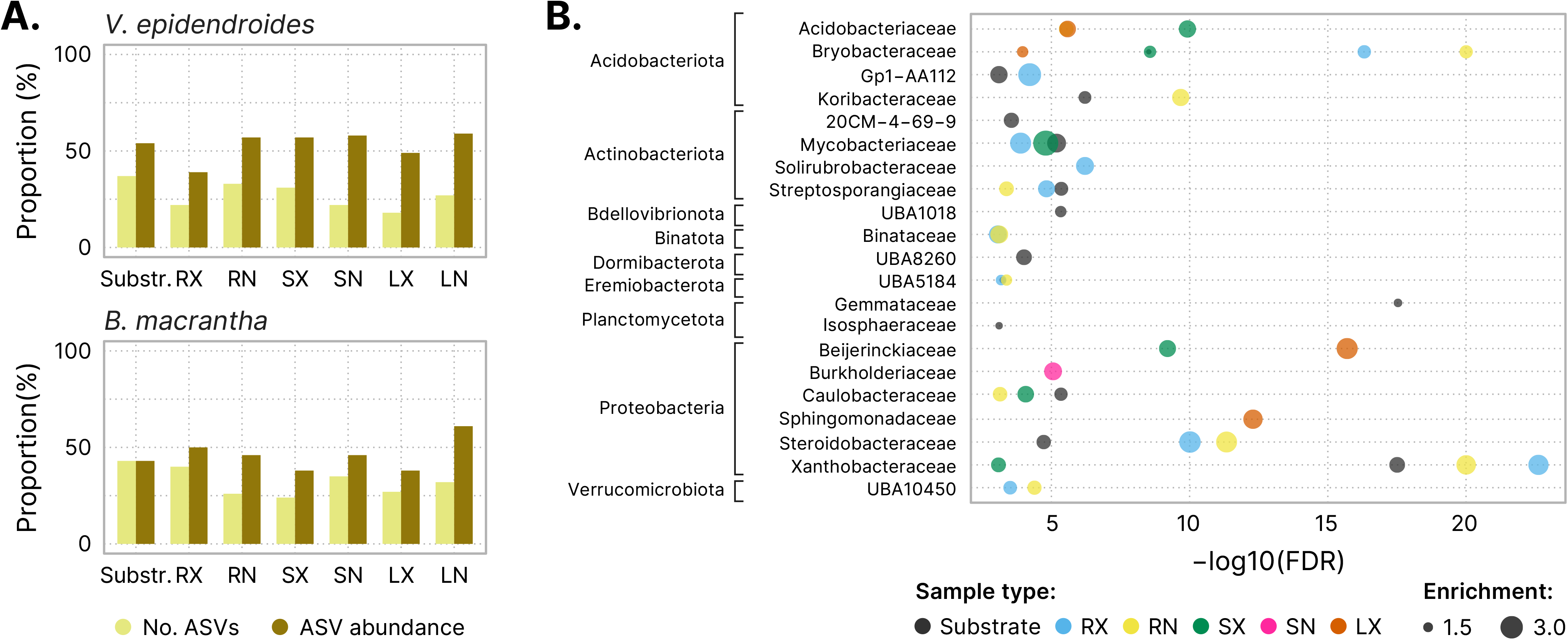
**(A)** Proportion of the total number of 16S ASVs (light yellow) and of the ASV abundance (dark yellow) shared between the communities associated with both plants. **(B)** Bacterial families (grouped by phyla) enriched within the shared ASV sets. The x-axis shows the false discovery rate (FDR) obtained from hypergeometric tests. The extent of the enrichment for each family, represented by the circle areas, was quantified as the ratio between the number of ASVs in the shared fraction and the number of ASVs observed in both the shared and exclusive fractions. No family was found to be enriched in the internal leaf communities of either plant. RX = root (external), RN = root (internal), SX = stem (external), SN = stem (internal), LX = leaf (external), LN = leaf (internal).

To determine whether shared bacterial species exhibited extensive genetic divergence between populations on each host plant, we selected MAGs that were highly abundant in both plants and evaluated differences in allele frequencies, nucleotide diversity, and linkage disequilibrium. We found multiple hallmarks of populational divergence that seemed to be tied to the genome taxonomy and that were likely shaped by non-neutral evolutionary processes [24] (Suppl. Note 3, Suppl. Figure 4).

### Evaluation of microbiome recruitment by the hosts and the microbial carbon cycling

Plants recruit beneficial microorganisms dynamically and selectively using root exudates rich in organic compounds such as amino acids and organic acids [21,25,26]. To investigate whether *V. epidendroides* and *B. macrantha* actively shape their microbiomes, the abundances of genes encoding organic substrate transporters (Suppl. Table 8) were estimated across all substrate and rhizosphere metagenomes. We found that the abundances of the evaluated transporters was systematically higher in the rhizosphere communities (Figure 4, LMM *p*-value < 0.001; ω^2^ ≈ 0.14). Transporter enrichment in the rhizosphere occurred for all organic substrates that were evaluated (Suppl. Table 9), suggesting that multiple molecules may be involved in the recruitment process. Although the two plant species did not significantly differ in total transporter abundance they exhibited differing taxonomic profiles (Suppl. Figure 5A).

**Figure 4.**
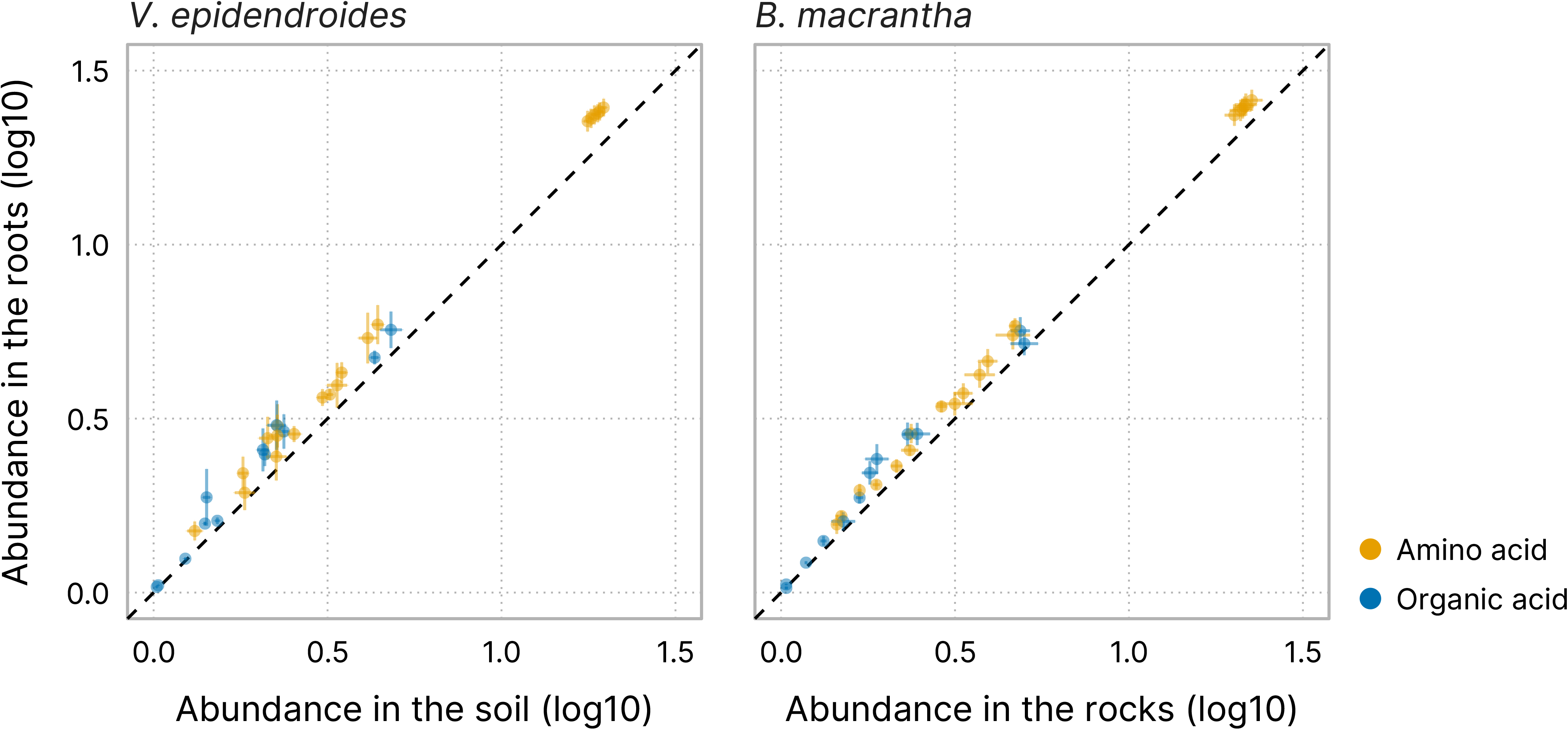
Mean total abundance (sum of RPKGs) of the investigated transporter genes in the substrates (x-axis) and roots (y-axis) of both plants. Circles are colored according to their assigned substrate class: amino acids (23 substrates) and organic acids (16 substrates). Horizontal and vertical lines represent the standard error of the mean in the substrates and roots, respectively.

Because the soil substrate contains higher organic matter content than the exposed rocks [27], we investigated the carbon cycling potential of the *V. epidendroides* and *B. macrantha* microbiomes with respect to carbohydrate degradation and carbon dioxide fixation (Suppl. Note 4, Suppl. Figure 5, Suppl. Tables 10 and 11). We found that genes associated with carbohydrate turnover were more abundant in the *B. macrantha*-associated communities (Suppl. Figure 5D). On the other hand, the microbiomes of both plants exhibited potential for autotrophy, likely through different mechanisms for energy generation as photosynthetic bacteria were much more abundant in the rock dwelling communities (Suppl. Figure 5E, F). Through metabolic potential inference we identified 38 autotrophic MAGs belonging to diverse lineages, including new families (Suppl. Table 11).

### Investigation of the P turnover potential by Velloziaceae-associated microbiomes

Microorganisms encode diverse mechanisms for P mobilization [28], and as their biomass turns over through time, P becomes available to plants [29] (Figure 5A). To investigate the P turnover potential of the *V. epidendroides* and *B. macrantha* microbiomes, we measured the total abundance of genes involved in environmental P mobilization (Suppl. Table 12). Multiple processes were highly represented in the metagenomes of both plants (Figure 5B) and were carried out by diverse taxa (Suppl. Figure 6A). Interestingly, some abundant P mobilization processes, such as the exopolyphosphatase activity and the catabolism of phosphanates, are not used by plants to increase P uptake, indicating a complementarity between repertoires of the hosts and their microbiome (Figure 5A). Additionally, a systematic increase in the abundances of P turnover processes in the rhizospheres relative to their adjacent substrates was verified (Figure 5B, LMM *p*-value < 0.001; ω^2^ ≈ 0.12).

**Figure 5.**
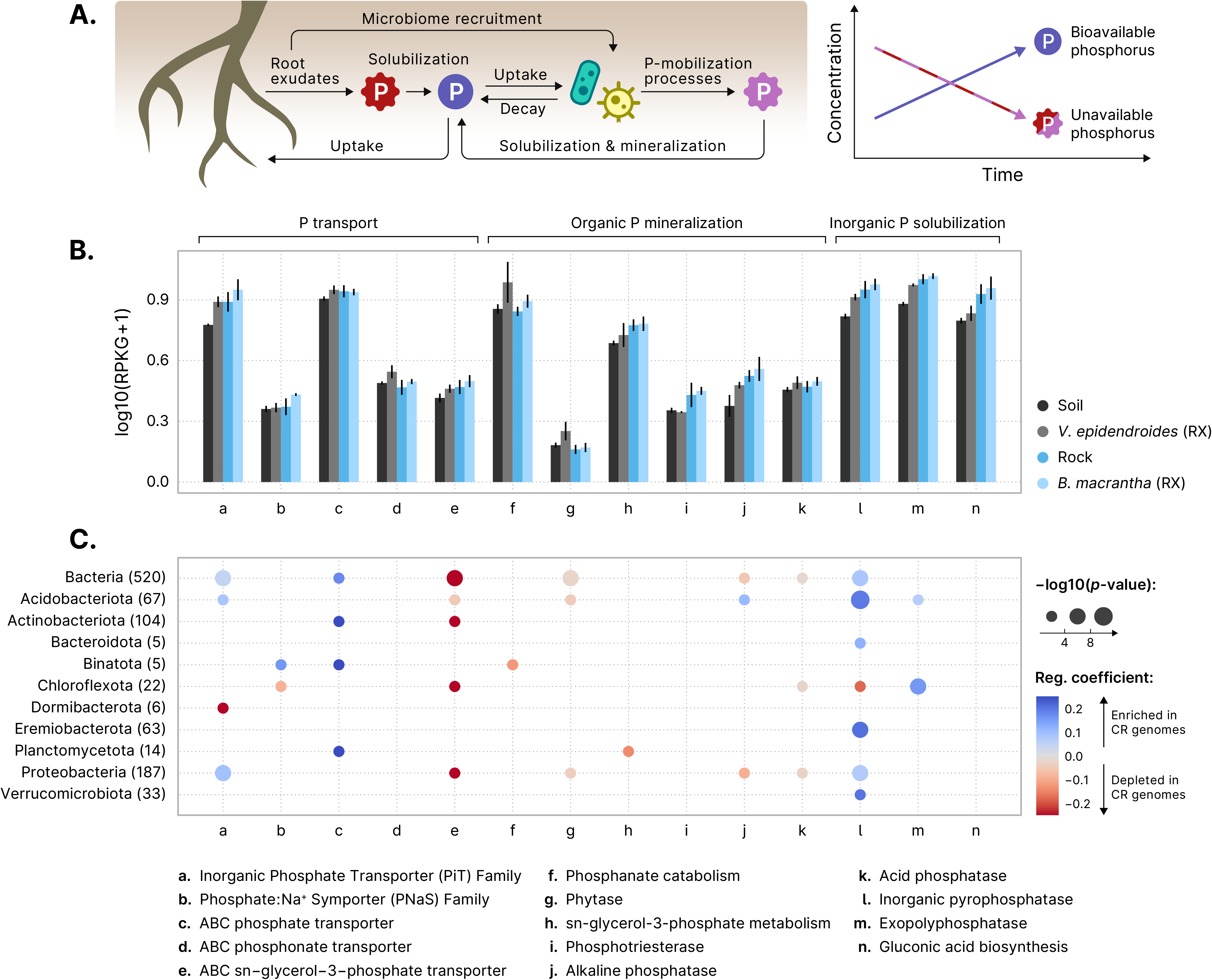
**(A)** Root exudates both solubilize phosphorus (P) in the plant substrate and recruit microorganisms that consume this nutrient. As the recruited microbes can mobilize phosphorus that would otherwise be unavailable for the plants (in pink), the total bioaccessible phosphorus concentration increases over the time. **(B)** Mean total abundances (sum of RPKGs) of proteins and pathways involved in processes linked to phosphate turnover (transport, mineralization, and solubilization) in the substrate and root-associate communities. Abundances of multiprotein complexes (*pstABCS*, *phnCDE*, *ugpABCE*, and *phnGHIJKLM*) were computed by averaging the abundances of their subunits. Vertical lines represent the standard error of the mean. **(C)** Phylogenetic regressions of the number of phosphate turnover-related genes. The MAGs retrieved in this study were compared to GTDB genomes to identify differences in the numbers of gene copies associated with phosphorus turnover. The color scale indicates the magnitude of the enrichment (blue) or depletion (red) of each process in the MAGs, and the area of the circles represent the statistical significance of the regression coefficient. Regression coefficients with *p*-value > 0.05 are omitted. Phylogenetic regressions were performed on the whole set of bacterial genomes and on the phyla containing at least 5 MAGs. RX = root (external), CR = *campos rupestres*.

Despite extensive microbiome specialization in the two plants, 12 out of 15 families with the highest total abundances of genes involved in P turnover were found to be significantly enriched in their shared microbiome fraction (Table 1, Figure 3B), indicating that the microbiome shared between these two plants played an important part in P nutrition. We also verified that different taxa might contribute to P mobilization using distinct mechanisms. For example, the Bryobacteraceae exhibited a high abundance of the *gcd* gene, involved in the synthesis of gluconic acid, a molecule that solubilizes P by chelating cations that are bound to recalcitrant phosphate [28, 30]. In contrast, the UBA5184 had high abundances of genes encoding exopolyphosphatase [31] and inorganic pyrophosphatase [32], enzymes that release P from inorganic phosphate polymers. Interestingly, many of the families with the highest P turnover potential had high PG, indicating that the novel lineages found in the *campos rupestres* play a part in P mobilization.

**Table 1.** The top 15 families with the highest average total abundance of processes associated with P turnover. Families that were significantly enriched among the AVSs that were shared between the two plants are indicated in bold text. The values under each sample type represent the average total RPKG of processes associated with phosphorus turnover. The rightmost column indicates the phylogenetic gain (PG) of each family. RX = root (external).

| Family | Phylum | *V. epidendroides* |  | *B. macrantha* |  | All samples | Family PG (%) |
| --- | --- | --- | --- | --- | --- | --- | --- |
|  |  | Soil | RX | Rock | RX |  |  |
| Undetermined | Undetermined | 27.52 | 32.25 | 27.39 | 28.79 | 28.99 | N/A |
| **Xanthobacteraceae** | **Proteobacteria** | **8.15** | **11.40** | **13.84** | **15.39** | **12.20** | **16.23** |
| **Bryobacteraceae** | **Acidobacteriota** | **2.54** | **3.03** | **6.30** | **7.83** | **4.92** | **32.15** |
| **UBA5184** | **Eremiobacterota** | **5.92** | **6.31** | **3.64** | **3.23** | **4.77** | **47.05** |
| Acetobacteraceae | Proteobacteria | 3.53 | 4.50 | 3.12 | 6.30 | 4.36 | 18.00 |
| **Solirubrobacteraceae** | **Actinobacteriota** | **0.83** | **2.01** | **6.60** | **6.39** | **3.96** | **34.32** |
| **Burkholderiaceae** | **Proteobacteria** | **3.77** | **7.47** | **1.71** | **2.00** | **3.74** | **2.01** |
| **Beijerinckiaceae** | **Proteobacteria** | **2.93** | **3.42** | **3.36** | **4.20** | **3.48** | **13.14** |
| Reyranellaceae | Proteobacteria | 2.60 | 2.28 | 1.45 | 0.88 | 1.80 | 34.50 |
| **Streptosporangiaceae** | **Actinobacteriota** | **1.23** | **1.12** | **1.94** | **1.84** | **1.53** | **6.43** |
| **Steroidobacteraceae** | **Proteobacteria** | **1.99** | **1.66** | **1.14** | **1.28** | **1.52** | **24.15** |
| **Acidobacteriaceae** | **Acidobacteriota** | **2.23** | **2.18** | **0.71** | **0.79** | **1.48** | **3.46** |
| **Sphingomonadaceae** | **Proteobacteria** | **0.31** | **0.73** | **2.13** | **2.52** | **1.42** | **2.94** |
| URHD0088 | Proteobacteria | 1.50 | 1.65 | 0.93 | 1.26 | 1.33 | 91.35 |
| **Mycobacteriaceae** | **Actinobacteriota** | **1.19** | **1.16** | **1.00** | **1.66** | **1.25** | **2.16** |
| **Caulobacteraceae** | **Proteobacteria** | **0.87** | **1.61** | **1.20** | **1.17** | **1.21** | **6.50** |

We compared the MAGs to genomes from GTDB r89 to evaluate whether they displayed trends regarding the number of genes involved in P turnover processes. Phylogenetic regression models revealed that multiple processes were significantly enriched in the *campos rupestres* MAGs, including two types of phosphate transporters (the PiT and the ABC transporter families), inorganic pyrophosphatases and exopolyphosphatases (Figure 5C). Interestingly, the MAGs showed a significant reduction in the number of genes involved in the transport of sn−glycerol−3−phosphate and in the hydrolysis of phytate, indicating that these compounds might not be reliable sources of P in the *campos rupestres*.

Siderophores, synthesized by biosynthetic gene clusters (BGCs) and primarily used to scavenge iron [33, 34], also take part in P turnover by releasing P bound to iron cations. The assembled metagenomes had 42 siderophore-producing BGCs, which were grouped according to their domain organization and sequence similarity into 15 gene cluster families (GCFs) (Figure 6). We found that a large gene cluster clan (GCC) — encompassing three GCFs and 18 siderophore BGCs (blue labels in Figure 6) and assigned to the Pseudonocardiaceae family — possessed domains similar to those of desferrioxamine siderophores [35, 36]. However this GCC had distinctive domain organization and in tandem duplication of the siderophore biosynthesis protein (IucA/IucC family), suggesting significant structural differences between their final products and described desferrioxamines. All siderophores within the Pseudonocardiaceae GCC were comparatively much more abundant in *B. macrantha* root and rock communities, which was supported by 16S data (Suppl. Figure 6B), and similar abundance profiles were observed across most of the other siderophore BGCs. This difference in the siderophore synthesis potential was further supported by the fact that the “Biosynthesis of siderophore group nonribosomal peptides” KEGG pathway was significantly enriched in the *B. macrantha*-associated communities (FDR < 0.001).

**Figure 6.**
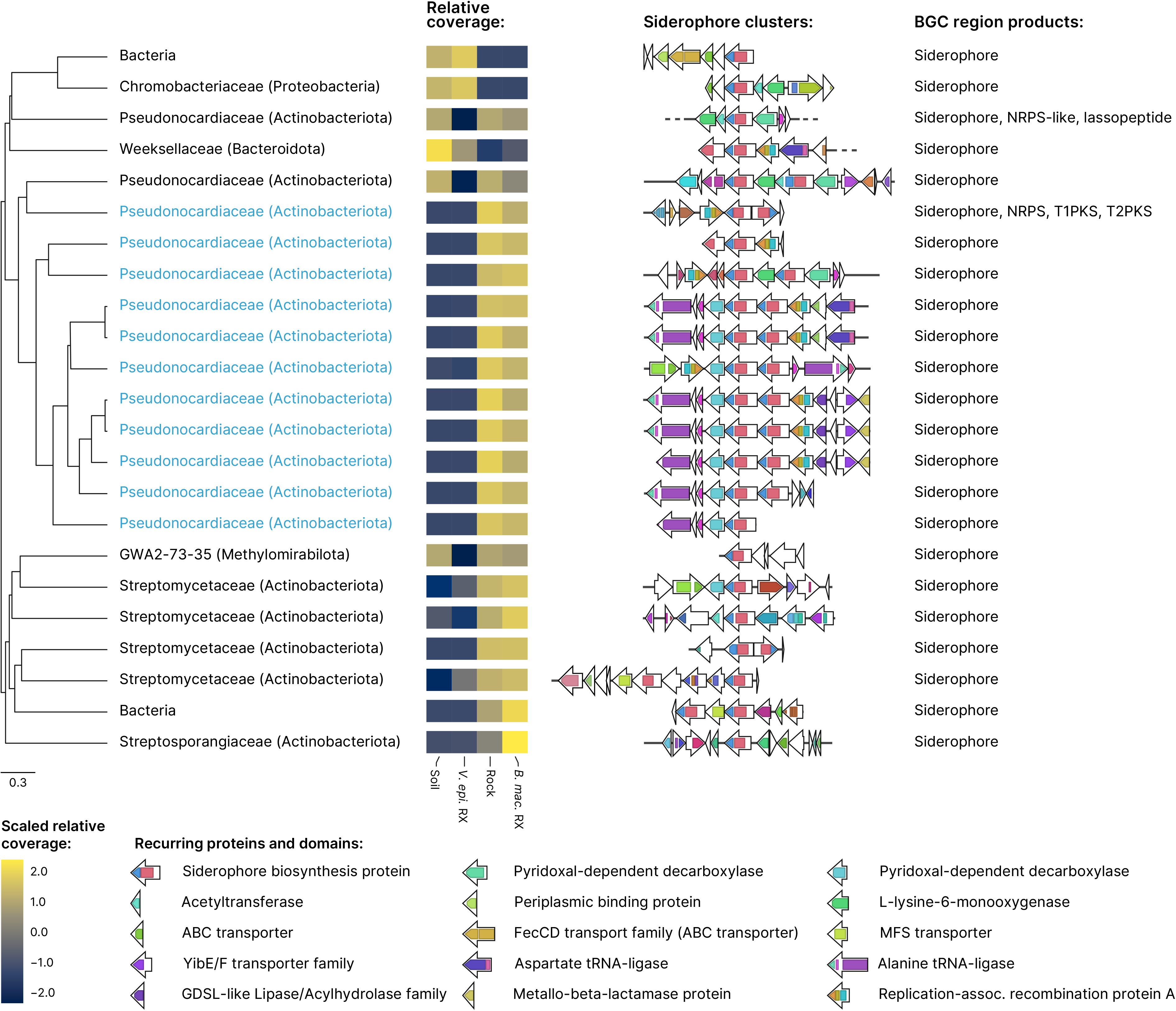
Structural diversity of siderophore biosynthetic gene clusters (BGCs) identified in the *campos rupestres* metagenomes. BGC regions containing siderophore clusters were hierarchically clustered using UPGMA with BiG-SCAPE distances. Groups of highly similar regions were identified based on their inconsistency coefficient and only the medoids are shown. Blue labels indicate the BGC regions that belong to the Pseudonocardiaceae-associated gene cluster clan. Taxonomies are presented at the family and phylum (in parenthesis) levels, except for two BGC regions whose contigs were assigned to the Bacteria domain. Heatmaps represent scaled means of log-transformed relative contig coverages in the four environments. Gene clusters are shown as arrays of genes (arrows) and their protein domains (colored blocks) centered at the siderophore biosynthesis protein. BGC regions containing other types of biosynthetic clusters (rightmost column) were trimmed to display only the loci assigned to the siderophore clusters.

Although arbuscular mycorrhiza are well known to increase P uptake by roots [37], Velloziaceae growing on severely P-impoverished substrates exhibit extremely reduced colonization by mycorrhizal fungi [7,15,38]. However, the ITS data showed that both plants harbored diverse endophytic fungi root communities (Suppl. Figure 3A, B), which compelled us to interrogate whether the fungi associated with *V. epidendroides* and *B. macrantha* participate in P nutrition. Using a read-level targeted gene finding approach, we surveyed the metagenomic data to identify orthologs of the fungal high-affinity H^+^:Pi transporter (*PHO84*), which participates in both P uptake from the substrate and phosphate efflux to the plant in the symbiotic interface [39], which, in at least one case, is required for the establishment of endosymbiosis [40]. In total, 312 fungal *PHO84* were retrieved, 67.6% of which were assigned to taxa containing known endophytes, such as the Dothideomycetes, Sordariomycetes, Leotiomycetes, Agaricomycotina, and Glomeromycetes [41–44]. Analysis of their combined abundance revealed that, as expected for symbiotic fungi, *PHO84* was enriched in the rhizospheres relative to the adjacent substrates, albeit with low statistical significance (Suppl. Figure 6C, LMM p-value ≈ 0.18, ω^2^ ≈ 0.1).

### Reconstruction of the N cycle dynamics in the *V. epidendroides* and *B. macrantha* microbiomes

Bacteria and Archaea contribute to the nitrogen (N) cycle by participating in several steps such as the release of N from organic matter, N fixation into ammonia, nitrification, and denitrification (Figure 7A). To understand how microorganisms mediate N turnover in the *campos rupestres* and how they can impact plant nutrition, we evaluated the taxonomic profile and the abundances of genes involved in different steps of the nitrogen cycle (Suppl. Table 13). Several taxa possessed the genomic potential to mediate nitrogen transformations (Figure 7A, Suppl. Figure 7A). Abundances of genes involved in cycling N were slightly higher in the rhizospheres of *V. epidendroides* and *B. macrantha* in comparison to their adjacent substrates (Figure 7B, LMM p-value ≈ 0.06, ω^2^ ≈ 0.02), suggesting that the potential to transform N is a feature of some root-recruited microorganisms.

**Figure 7.**
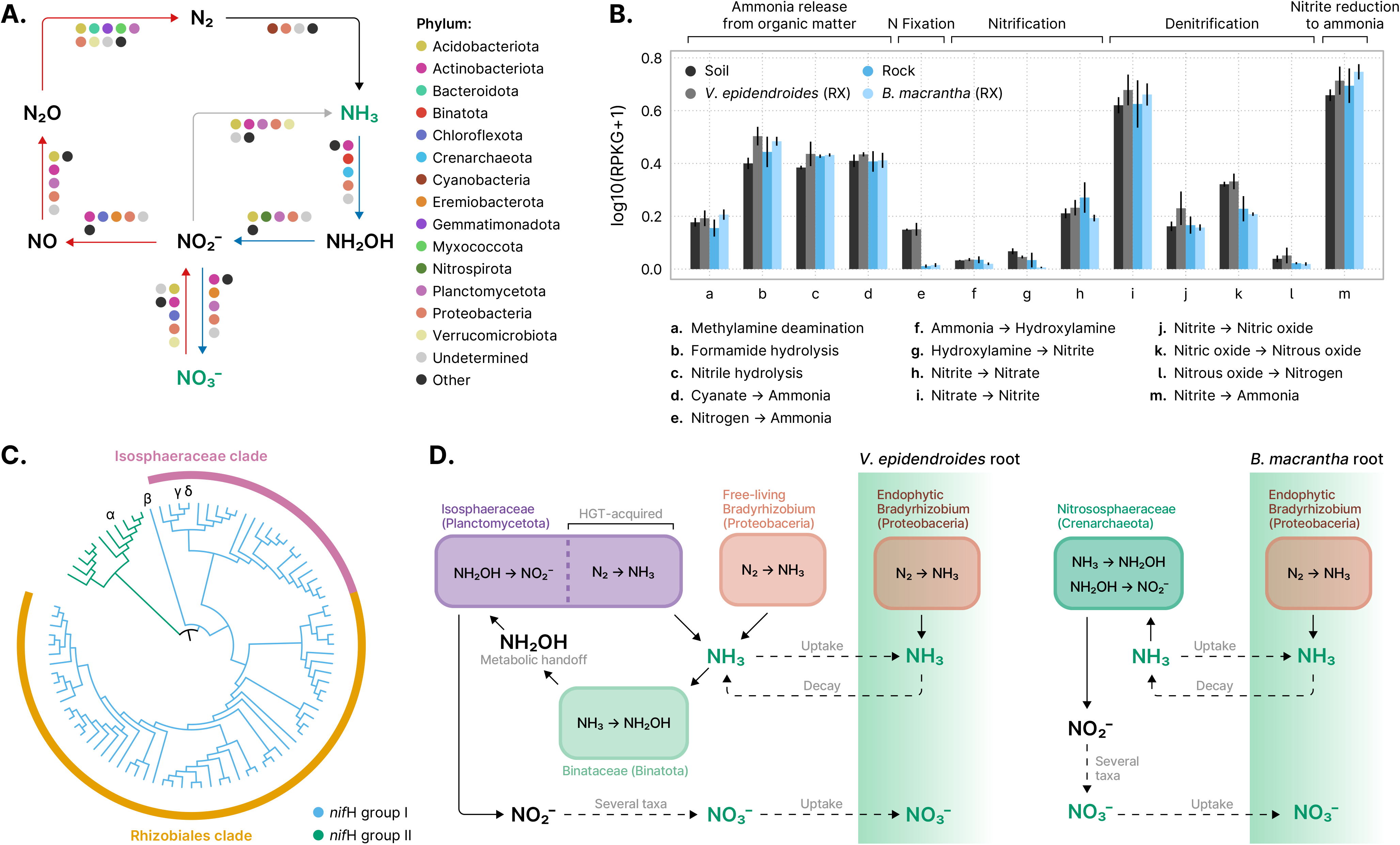
**(A)** Phyla predicted to be involved in nitrogen-cycling reactions: fixation (black arrow), nitrification (blue arrows), denitrification (red arrows), and nitrite reduction to ammonia (gray arrow). Compounds that can be taken up by plants roots are depicted in green. Taxonomic assignment was performed at the contig level. Phyla that contributed less than 5% of the detected genes involved in each reaction were grouped under the “Other” category. **(B)** Total abundances (sum of RPKGs) of reactions involved in nitrogen turnover processes in the substrate and external root-associated communities. The abundances of multiprotein complexes (*nifHDK*, *amoABC*, *nxrAB*, *narGHI*, *napAB*, *nasAB*, *norBC*, *nirBD*, and *nrfAH*) were computed by averaging the abundances of their subunits. Vertical lines represent the standard error of the mean. RX = root (external). **(C)** Cladogram of maximum-likelihood phylogenies inferred from a dereplicated set of metagenomic *nifH* orthologs. Branches are colored according to the major *nifH* group they belong to, and the tree was rooted in the node between the two groups. The dominant clades are indicated by the outer rings. Orthologs encoded by MAGs containing at least two *nif* subunits are indicated by greek letters: (α): Verrucomicrobiota, (β): Enterobacteriaceae, (γ and δ): Isosphaeraceae. **(D)** In the *V. epidendroides*-associated communities, nitrogen is mostly fixed by endophytic and free-living *Bradyrhizobium*, and by Isosphaeraceae, which most likely received their *nif* complex via horizontal gene transfer (HGT). The produced ammonium is then oxidized into hydroxylamine by methylotrophic Binataceae. This molecule is then released from the cell and is oxidized into nitrite by Isosphaeraceae. In the *B. macrantha*-associated communities, nitrogen is likely converted into ammonium by endophytic *Bradyrhizobium*. Ammonia originated from organic matter decomposition is oxidized into hydroxylamine and, subsequently, into nitrite by Nitrososphaeraceae. In both plants, the oxidation of nitrite into nitrate is likely performed by a several taxa.

We set out to further survey the N fixation profile of the *campos rupestres* communities by probing the metagenomes and MAGs for genes encoding subunits of the nitrogenase complex (*nifHDK*). Sequence-based taxonomic assignment of nitrogenase subunits and phylogenetic analysis of the dereplicated set of all *nifH* subunits revealed that most of the *nif* diversity comes from the Rhizobiales order (Proteobacteria phylum) and the Isosphaeraceae family (Planctomycetota phylum) (Figure 7C, Suppl. Figure 7B). We also retrieved *nifH*-containing MAGs assigned to the Enterobacteriaceae family and Verrucomicrobiota phylum, the latter of which encoded a *nifH* from the group II clade, that was predominantly comprised of orthologs from obligate anaerobes [45].

Even though there is no report of an Isosphaeraceae-encoded *nif*, we found all subunits of the complex in four MAGs belonging to two species clusters, which persuaded us to reconstruct the evolutionary history of these orthologs. A phylogenetic analysis using *nifH* and *nifD* sequences from the Isosphaeraceae MAGs, other Planctomycetota species, and orthologs with high sequence similarity revealed that nitrogenase encoded by the MAGs formed a clade with Gammaproteobacteria, rather than Planctomycetota, strongly suggesting that they acquired *nifHDK* via horizontal gene transfer (HGT) (Suppl. Figure 7C). Indeed, we found clear synteny between a *nif*-containing Isosphaeraceae contig and the *Pseudomonas stutzeri* genome, which harbors a packed *nif* cluster [46]. Additionally, gene-level taxonomic assignment revealed clear boundaries between the genes vertically inherited from ancestral Isosphaeraceae and genes within the region received via HGT (Suppl. Figure 7D).

As for the Rhizobiales, the vast majority (≈ 91.5%, or 54/59) of the orthologs prior to dereplication were found in unbinned contigs from *V. epidendroides* and soil metagenomes, resulting in an apparent major difference between the N fixation potential of the communities associated with the two plants. As the metagenomic data mostly captured free-living populations and because endophytic bacteria are major players in the N fixation process, we examined the abundances of *Bradyrhizobium* — a genus within the Rhizobiales that encompasses several endophytic diazotrophs — using 16S data and found that these bacteria were enriched in the endophytic compartments of both plants (Suppl. Figure 7E). To evaluate whether these endophytes can fix N, we used sensitive read-level gene identification to probe *Bradyrhizobium nifH* sequences in metagenomic data, which revealed that *nifH* assigned to *Bradyrhizobium* were present in the *B. machantha* and rock metagenomes, albeit in lower levels than in the *V. epidendroides*-associated communities, and that they were enriched in the rhizospheres (Suppl. Figure 7F, LMM *p*-value < 0.05; ω^2^ ≈ 0.45). We also identified two *Bradyrhizobium* contigs, retrieved from the metagenomes of both plants, that harbored proviral sequences containing the *exoZ* gene, which encodes a cell surface modifying acetylase that increases the efficiency of endosymbiosis establishment [47] (Suppl. Figure 7G). Comparison with IMG/VR’s [48] viral genomes revealed that similar bacteriophages of the Myoviridae family were previously detected in root nodule and rhizosphere communities.

Besides the ammonia generated by N fixation, the primary source of inorganic nitrogen to plants is nitrate, which is produced through sequential oxidation of ammonia (Figure 7A). These reactions are carried out by ammonia-oxidizing Bacteria (AOB) and Archaea (AOA), which form nitrite in two steps (converting ammonia into hydroxylamine, and hydroxylamine into nitrite), and by nitrite-oxidizing bacteria, which convert nitrite into nitrate (Figure 7A). Inspection of the taxonomic assignments of contigs containing ammonia monooxygenase (*amoABC*) and hydroxylamine oxidoreductase (*hao*) revealed that the Proteobacteria, which encompass all the traditional AOB [49], are a minor fraction of the nitrifying communities of the *campos rupestres* (Suppl. Figure 7A).

Canonically, both steps of nitrite formation are carried out by bacteria harboring both *amoABC* and *hao*; however, we found that the taxonomic profiles of these two enzymes were surprisingly contrasting, suggesting that these reactions might be performed by distinct populations (Suppl. Figure 7A). Inspection of the MAGs’ gene repertoires supported this hypothesis, as no genome encoded both the *amo* complex and *hao* (Suppl. Figure 7H). Two groups of ammonia-oxidizing MAGs were retrieved: the Nitrososphaeraceae, a family of AOA that harbors *amoABC* and is known to oxidize hydroxylamine into nitrite without *hao* [50]; and the Binataceae, a family of the methylotrophic phylum Binatota which encodes the methane monooxygenase cluster (*pmoABC*), a close ortholog of *amoABC* that has been shown to oxidize ammonia [51]. As for *hao*, we found highly abundant Isosphaeraceae MAGs encoding orthologs that contained all the 16 heme-binding cysteines necessary for function. Curiously, all the N-fixing Isosphaeraceae also contained *hao*, suggesting that these genomes fulfill a dual role in the nitrogen cycle. Furthermore, Isosphaeraceae genomes in GTDB r89 contained *hao*, indicating that ancestral lineages already participated in N turnover.

## Discussion

Our data show that the microbiomes associated with the endemic *V. epidendroides* and *B. macrantha* are diverse and our results greatly expand the known diversity of several bacterial lineages. Notably, we provide evidence that these microorganisms are likely recruited by the plants and play a role in nutrient turnover, likely contributing to plant fitness.

### The *campos rupestres* microbial communities harbor a high degree of taxonomic and function novelty

At high ranks, the taxonomic profiles of the *campos rupestres* microbiomes echoed that of global soil surveys for bacteria, archaea, and fungi [19–21], with communities dominated by phyla such as Proteobacteria, Acidobacteriota, Verrucomicrobiota, Ascomycota and Basidiomycota (Figure 1B, Suppl. Figure 1A). However, as the *campos rupestres* constitute an underexplored biodiversity hotspot with high levels of plant endemism, we evaluated the uniqueness of the communities associated with *V. epidendroides* and *B. macrantha* using multiple approaches and discovered extensive taxonomic and functional novelty.

Our data expanded the phylogenetic diversity of multiple understudied groups (Figure 1E), including the Eremiobacterota, that are involved in phosphorus turnover (Table 1) and carbon fixation (Suppl. Figure 5E, F), the Dormibacterota, also involved in carbon fixation (Suppl. Figure 5F), and the Binatota, a group of methylotrophic bacteria [52] that likely participates in nitrogen cycling in the *campos rupestres* soils (Figure 7D, Suppl. Figure 7A, H). Additionally, we found that the below-ground fungal communities were significantly more novel than those colonizing the stem and leaves.

Regarding the functional novelty, a drastic reduction of the fraction of annotated clusters was observed when proteins were grouped at 50% sequence identity, suggesting that the microbial communities of the *campos rupestres* harbor several small and lineage-specific gene families. This finding was further supported by comparative genomic analysis, which revealed hundreds of orthogroups that were exclusive to the genomes assembled in this work. We note, however, that because most of the existing bacterial diversity has not been sequenced the number of novel orthogroups is likely being overestimated.

The microbiomes of *V. epidendroides* and *B. macrantha* are highly differentiated but share a core of highly efficient colonizers

Even though the rocks over which *B. macrantha* grows are nutrient depleted as compared to the soils — especially regarding organic matter and nitrogen — we found that the prokaryotic diversity did not significantly differ between the microbiomes of the two plants, suggesting that the rock-dwelling microorganisms possess mechanisms that allow them to grow under severe limitations. Indeed, we found evidence that phosphorus and nitrogen turnover were not reduced in the rocks. In contrast, fungal communities associated with *B. macrantha* were less diverse than the ones associated with *V.epidendroides*, which is in accordance with previous reports that plants growing over rocks have reduced fungal colonization in the *campos rupestres* [7].

The different microbiomes of the two plants, despite their geographic proximity [27], suggests that contrasting substrates and host biology shape plant-associated communities in the *campos rupestres*. Furthermore, the finding that particular taxa become specialized on their host indicates that host and environment-specific pressures select specific functions. This is substantiated by the diverging genic repertoire of the *V. epidendroides* and *B. macrantha* microbiomes, with abundances of multiple metabolic pathways differing between them. One notable example of such structured differentiation is the recurrent enrichment of Actinobacteriota families in *B. macrantha*, which can be attributed to the resilience of these bacteria to low humidity [53] and greatly influenced the siderophore production in the rock-dwelling communities.

Despite microbiome specialization, the two plants shared a core microbiome of efficient microbial colonizers that were likely adapted to the harsh environmental conditions of the *campos rupestres*. Accordingly, several of the bacterial families enriched among the shared ASVs (Figure 3B) were involved in phosphorus turnover, suggesting that these core taxa may increase plant fitness and were likely recruited by both plants.

### The Velloziaceae microbiomes likely contribute to phosphorus availability and are recruited by root exudates

Species of the Velloziaceae family increase P availability in soils [15] and rocks [5] by secreting exudates containing carboxylates that release phosphate bound to cations, such as iron and aluminum [54]. Despite these adaptations, some forms of P are unavailable for plant uptake and are only accessed by bacteria, which harbor diverse processes that make P bioaccessible [28]. As root exudates are rich in organic compounds that recruit and sustain microbial communities [21], we hypothesized that Velloziaceae exudates fulfill a dual role: they increase the labile P concentration by direct solubilization and also recruit microorganisms that possess a complementary repertoire of molecular processes that increase P availability in the root’s vicinity. In this scenario, microorganisms would use phosphate for their own needs, but would benefit the system in the long term by mobilizing P otherwise unavailable to plants [29] (Figure 5A).

We assessed active recruitment of microorganisms by evaluating the abundances of transporters with specificity for amino acids and organics acids that are secreted by roots [25, 26]. The increased abundances of these transporters in the rhizosphere communities of both plants (Figure 4) suggests that microorganisms able to take up organic compounds were being selected in the vicinity of the roots, which is consistent with our hypothesis that Velloziaceae root exudates shape the plant microbiome. The enrichment of processes linked to P turnover in the root microbiomes compared to the substrates indicates that the recruited bacteria can increase bioaccessible P (Figure 5B). Furthermore, the fact that most of the taxa with elevated P turnover potential were enriched within the shared microbiome of *V. epidendroides* and *B. macrantha* (Table 1) suggests that both plants actively recruit a common set of nutritionally relevant microorganisms. To illustrate this scenario, we show that the Bryobacteraceae, whose known members are acidophilic chemoheterotrophs that consume organic acids [55], were enriched within the shared microbiome, had high abundances of genes involved in P turnover, and had high abundances in rhizospheres (Table 1).

By evaluating the production of siderophores by plant-associated bacteria, we found that the examined communities harbored structurally diverse BGCs, including some that form hybrid regions with other classes of biosynthetic clusters (Figure 6, rightmost column), which suggests a diversified repertoire of final siderophore products. In addition, the siderophore repertoires of *B. macrantha*-associated communities were much larger, which could be interpreted as an indirect consequence of the higher abundance of the BGC-rich [56] Actinobacteriota in these microbiomes. We believe, however, that a complementary nutrient-driven explanation is also appropriate: as the iron content of the sampled rocks is around 8-fold lower than that of the soils [27] this nutrient is under increased demand in *B. macrantha* microbiomes, increasing the fitness of Actinobacteriota populations that are efficient at scavenging iron and, consequently, solubilizing phosphate.

Despite previous reports suggesting that Velloziaceae growing under severe P-impoverishment exhibit reduced fungi colonization [7,15,38], a diverse set of transporters were assigned to fungal taxa that comprise known endosymbiotic species and that were enriched in the rhizosphere (Suppl. Figure 6C). Given that the below-ground fungal communities under investigation included highly novel fungal lineages (Suppl. Figure 1B), we hypothesize that undescribed species may establish associations with the radicular systems of both plants, occupying the ecological niche left by arbuscular mycorrhiza whose abundances were extremely reduced in the *campos rupestres* (median Glomeromycota ASV abundance in below-ground samples: 0.23% and 0.20% in *V. epidendroides* and *B. macranta*, respectively).

### Nitrogen cycling is heavily influenced by the substrate and involves newly described microbial lineages and interaction dynamics

In the *campos rupestres*, nitrogen is constantly lost from the biological pool due to seasonal fires [1]. Therefore, as biological fixation of atmospheric N is likely pivotal to sustain the biomass of this ecosystem, we set out to investigate the N cycle in the context of the *V. epidendroides* and *B. macrantha*-associated communities. Our finding that N-transforming processes in the *campos rupestres* were carried out by several taxa challenges the traditional view that some reactions, such as ammonia and nitrite oxidation, are carried out by few restricted clades. Additionally, as we observed for genes involved in P-turnover, we found that N cycle processes were also enriched in the rhizosphere, although at a much lower level, suggesting that microorganisms that can perform some of the transformations in the N cycle have increased fitness when associated with plants, possibly due to active root recruitment.

We identified multiple Isosphaeraceae genomes with metabolic potential for nitrogen fixation. Although *nifHDK* genes have been documented in some marine Planctomycetota [57], their complexes belong to the group II *nif*, while the *campos rupestres* Isosphaeraceae orthologs belong to group I (Suppl. Figure 7B, C). A combination of approaches indicates that the N fixing potential of the Isosphaeraceae MAGs was most likely the result of a HGT from a Gammaproteobacteria (Suppl. Figure 7C, D). We also speculate that the Isosphaeraceae, as other Planctomycetota, display cell compartmentalization [58], which could be used to create an anaerobic environment for *nif* activity, as observed in Planctomycetota possessing anammoxosomes [59]. Given that these MAGs compose a large fraction of the *campos rupestres nifH* diversity (Figure 7C, Suppl. Figure 7B) and that there is prior evidence for *nif* HGT events [46,60–62], we argue that horizontal transmission of the nitrogenase activity can affect the biogeochemical N cycle and, consequently, plant nutrition.

We also found a large amount of *nifH* orthologs that were assigned to the Rhizobiales order, a lineage that comprised several known diazotrophs. Interestingly, only a few of these orthologs were retrieved from *B. macrantha* and rock metagenomes and, although several *Bradyrhizobium* MAGs were recovered from these environments, none had the metabolic potential to fix N. We believe that the lack of *nif* is an advantageous trait for rock-dwelling bacteria, as the N fixation reaction is very carbon-demanding [63] and could cripple their growth in a carbon-poor environment [27]. We found, however, indirect evidence for endophytic diazotrophs as read-level gene identification revealed an enrichment of *Bradyrhizobium nifH* abundance in the rhizosphere of both plants (Suppl. Figure 7F). In addition, the finding that virus-encoded *exoZ* is integrated into *Bradyrhizobium* genomes (Suppl. Figure 7G) further supports that these bacteria establish endophytic associations, as phages employ auxiliary metabolic genes to enhance host fitness and boost their replication rates [64–66].

Regarding nitrification, we found no evidence for the participation of canonical AOB that possess the molecular machinery for the two-step ammonia conversion into nitrite within a single cell. Instead, we found that the ammonia and hydroxylamine oxidation reactions in the soil are likely carried out by distinct populations belonging to the Binataceae and Isosphaeraceae families, respectively. Even though *hao* has no other known function aside from hydroxylamine oxidation, *amoABC*/*pmoABC* can oxidize both ammonia and methane. In fact, some methylotrophs use *hao* as a hydroxylamine detoxification mechanism [51]. However, several Binatota genomes (22 genomes in GTDB r89 and 86 genomes in GEM [67]) encode *hao*, even though some encode genes involved in denitrification [52], which lead us to hypothesize that the methylotrophic Binataceae use metabolic handoff associations [68] with *hao*-encoding Isosphaeraceae as a means of hydroxylamine detoxification, benefiting the recipient cell by providing them with an energy-generating molecule.

As we investigated a broader genomic dataset, we found that the decoupling of the *amoABC*-*hao* system was commonplace, as approximately half (92 out of 193) of *hao*-containing bacterial genomes in GTDB (release 89, AnnoTree [69] annotation) have no subunit of the *amoABC* complex, including 16 Planctomycetota and 9 Acidobacteriota genomes (both phyla encoding *hao* in our metagenomes). Further strengthening our hypothesis, the alternative mechanism of hydroxylamine detoxification using the cytochrome P460 [70] was also absent in all Binatota MAGs and was found in 6 of 7 Isosphaeraceae genomes. We note, however, that even though hydroxylamine is known to be released from the cell to the environment [71, 72], it is a very reactive molecule, being readily oxidized into nitrogen trace gases [73, 74]. Therefore, close physical proximity is likely necessary for the metabolic handoff to be effective.

Altogether, our results suggest that most of the N fixation in *V. epidendroides* is performed by Isosphaeraceae and *Bradyrhizobium*, both free-living and endophytic. In *B. macrantha*, most fixation is likely performed by endophytic *Bradyrhizobium*, probably due to nutritional limitations of the substrate where this plant thrives (Figure 7D). The importance of *Bradyrhizobium* for plants is supported by the fact that its family (Xanthobacteriaceae) had the highest level of enrichment among the ASVs that were shared between the root communities (Figure 3B). As for nitrification, our MAG data suggest that ammonia oxidation is primarily conducted by Binataceae (in *V. epidendroides*) and by Nitrososphaeraceae (in *B. macrantha*), with a critical metabolic handoff interaction between Binataceae and Isosphaeraceae to convert hydroxylamine into nitrite (Figure 7D). Although many other taxa, such as Acidobacteriota and Proteobacteria, also may be involved in these reactions, we found no evidence for the participation of canonical AOB.

## Conclusions

This study set out to characterize the composition and functions of the microbiomes associated with two plant species that grow in the severely nutrient-impoverished substrates of the Brazilian *campos rupestres*. Our results indicate that microorganisms form close associations with their hosts and are involved in carbon, phosphorus, and nitrogen turnover. These findings highlight that plant-associated microorganisms contribute to plant nutrition and that assessing microbial diversity is crucial to understand the dynamics of nutrient cycling. We propose that future research of nutrient-limited environments considers the microbial diversity and ecology to develop holistic models of plant fitness.

## Methods

### Sample collection and metagenomic sequencing

The study design, sampling methodology, library preparation, and sequencing that were used in this study were thoroughly detailed previously [27]. Briefly, substrate (soil and rock), root, steam and leaves were sample from six individuals of *V. epidendroides* and *B. macrantha* in March of 2017, at the end of the wet season. For each plant species, a sampling area of approximately 200 m^2^ was defined and individuals with similar height, number of leaves, and number of tillers were selected and assigned random identifiers from R1 to R6. For *V. epidendroides*, the soil surrounding the plant within a 20 cm radius was excavated to a depth of 15 cm. For *B. macrantha*, the adjacent rocks were fragmented with a hammer and chisel until roots were exposed and the pieces of rock were collected and further grinded to small pieces. Host-associated microbiomes were gathered from the external and internal compartments of the root, steam, and leaves of the sampled plants using methods adapted from a previously described protocol [75].

Environmental DNA was extracted using the PowerSoil DNA Isolation kit (MO BIO Laboratories, Inc., Carlsbad, CA, USA). Amplicon sequencing of the 16S rRNA gene (V4 region) and the internal transcribed spacer 2 (ITS2), for profiling prokaryotic and fungal communities, was achieved through PCR amplification of the DNA extracted from all samples (substrates and the external and internal plant organs) using the 515FB/806R [76] and ITS9_Fwd/ ITS4_Rev [77] primer pairs and subsequent sequencing using the Illumina MiSeq platform to generate 2 × 300 bp reads. Metagenomic libraries of the external root and substrate communities from three individuals (samples R1 to R3) were generated with the Illumina HiSeq sequencing platform, yielding 2 × 150 bp reads. All library preparation and sequencing procedures were conducted at the DOE Joint Genome Institute.

### Amplicon sequence variants inference and taxonomic assignment

Sequencing reads of 16S and ITS were processed to remove the PCR primer sequences with cutadapt [78] (version 1.16). We retained read pairs where complete sequences of both the forward primer in the R1 read and the reverse primer in the R2 read were detected. Amplicon sequence variants (ASV) inference was then performed separately for the 16S and ITS libraries using DADA2 [79] (version 1.6.0). For the 16S libraries, we used Phred score profiles of the R1 and R2 reads to determine the lengths that the reads should be truncated to so that bad-quality regions were removed (245 bp for R1 and 180bp for R2). ITS reads were not truncated to a fixed length to facilitate overlap between the read pairs as this region presents significant length variation across genomes. Next, reads with ambiguous bases or with an expected number of errors greater than two were filtered out and DADA2’s error models were fitted to the R1 and R2 reads of both types of amplicon. These models and the dereplicated reads pooled from all samples were used as input for the core ‘dada’ function, which infers sequencing errors and correct read sequences. Finally, ASVs were obtained by merging read pairs whose R1 and R2 had an overlapped of at least 16 bp and by removing putative PCR chimeras.

Taxonomic assignment of ASVs was performed with the IDTAXA algorithm [80] (part of the DECIPHER library, version 2.8.1) using a minimum confidence threshold of 40% and the GTDB [81] (release 89) and UNITE [82] (Feb. 2020 release) databases as taxonomic references for 16S and ITS sequences, respectively. IDTAXA was also employed to identify ASVs derived from organellar genomes by assigning 16S sequences to taxa from the SILVA database [83] (release 138), which includes ribosomal genes of mitochondria and chloroplast. ASVs found to be derived from organellar genomes were excluded from downstream steps.

An additional filter was performed to remove likely spurious ASVs in an unsupervised manner, based on their recurrence across different samples [84]. We first computed the recurrence of each ASV as the number of samples in which it was observed at least once (that is, had at least one read associated to it) and then discarded ASVs with prevalence less than 2 were excluded from further analyses.

### Investigation and statistical analyses of ASV data

Community alpha diversity, quantified as both richness and evenness, was independently computed from 16S and ITS ASV count using the vegan library (version 2.5-5). Richness was estimated from count matrices rarified to 5,000 reads, a value that is close to the lowest sequencing depth among the samples. Community evenness was assessed through the Pielou’s equitability index. To test for relationships between the host plant species and community alpha diversity, the data was modeled using linear mixed-effects models (LMMs) with the lmerTest package (version 3.1-0) as follows: *Y* = *β* × *H* + S + C + E, where *β* is the regression coefficient, *H* encodes the host species, *S* and *C* are the random effects of the sampled individual and the sample type, respectively, and *E* is a vector of errors. The sample types used for modeling were the external and internal communities of the root, stem, and leaves; substrate samples were not included.

Beta diversity was appraised using Bray–Curtis and weighted UniFrac dissimilarities computed from relative ASV abundances using the phyloseq library [85] (version 1.34.0). The phylogenetic trees used to compute weighted UniFrac were reconstructed using IQ-TREE [86] (version 2.0.3, parameters: ‘-fast -m GTR+G4’) from 16S and ITS ASV alignments generated with MAFFT [87] (version 7.464, parameters: ‘--auto’). The statistical significance of grouping communities according to their host plant within the Bray–Curtis and UniFrac spaces was estimated using PERMANOVA (‘adonis function from vegan) [88].

The differentiation of communities belonging to the same type (substrate and the external/internal communities of the root, stem, and leaves) but associated with different host plants was appraised using two approaches. First, the main taxa driving the differentiation between *V. epidendroides* and *B. macrantha*-associated communities were identified with ALDEx2 [89] (version 1.18.0) with the parameters: denom = “zero”, mc.samples = 1000. The hypeR [90] library (version 1.6.0) was then used to identify statistically significant (FDR ≤ 10^−5^ and score ≥ 0.2) family-level taxa enrichment in each host plant by performing Kolmogorov-Smirnov tests on ASV lists ranked by their estimated effect size. The second approach to quantify community differentiation was the determination of the relative amount of shared ASVs between pairs of communities. Shared ASVs were defined as ASVs that had prevalence ≥ 2 within the samples of both communities in pairwise comparisons. Family-level taxa enrichment in the shared ASV sets was detected using hypergeometric tests (FDR ≤ 10^−3^), as implemented in hypeR.

To measure the taxonomic novelty of each community we devised the weighted average community identity (WACI) metric. Specifically, ASV sequences were aligned to reference databases (GTDB and UNITE for 16S and ITS sequences, respectively) using BLAST [91] (version 2.9.0) and community novelty was computed as the average ASV best hit alignment identity weighted by the relative ASV abundance in the sample. Differences between the WACI of below-ground (substrate and root-associated) and above-ground (stem and leaves-associated) communities were evaluated using LMMs as follows: *Y* = *β* × *R* + S + C + E, where *β* is the regression coefficient, *R* encodes the region (above or below ground), *S* and *C* are the random effects of the sampled individual and the sample type, respectively, and *E* is a vector of errors.

### Metagenome assembly and retrieval of metagenome-assembled genomes

Metagenome sequencing reads were quality-trimmed using cutadapt [78] (version 1.16) with the ‘--pair-filter=any -m 25 -q 5,5’ parameters, discarding read pairs where at least one of the reads was shorter than 25 bp after the trimming. To recover unpaired reads that were discarded because their pairs did not satisfy the length threshold, cutadapt was executed again on these pairs using only the ‘-m 25’ parameter. Both the trimmed read pairs and the unpaired reads were used for metagenomic assembly with using MEGAHIT [92] (version 1.2.7) with the ‘--k-min 27 --k-max 147 --k-step 10’ parameters. Additionally, four co-assemblies were generated by combining the reads from all samples of each sample type (soil, rock, and the rhizospheres of each plant) and assembling them together. Contigs shorter than 500 bp and 1,000 bp were filtered out from the individual and co-assemblies, respectively. Taxonomic assignment of the resulting contigs was attained with MAGpurify2 (available at https://github.com/apcamargo/magpurify2), using a database based on GTDB release 89 (doi: 10.5281/zenodo.3817702). We also performed protein-level assembly with PLASS (version 2.c7e35) [17], taking the trimmed read pairs as input. Only peptides containing both start and stop codons and with at least 60 amino acids were retained. This approach allowed us to recover a significant fraction of the protein sequence space that could not be attained by using the metagenomes alone (Suppl. Table 4).

Metagenome-assembled genomes (MAGs) were obtained by binning the contigs of each of the resulting assemblies and then dereplicating the bin sets recovered from assemblies of the same environment by clustering near-identical genomes. First, the coverage profile of contigs longer than 2 kb across all conditions was obtained using Bowtie 2 [93] (version 2.3.5) to cross-map the reads of each sample to each assembly. Next, the read mappings were used to bin each assembly using four different binning algorithms: MetaBAT2 [94] (version 2.14), MaxBin2 [95] (version 2.2.7), CONCOCT [96] (version 1.1.0), and Vamb [97] (version 3.0.1). DAS Tool [98] (version 1.1.2) was then executed to aggregate the four sets of bins generated from each assembly into 16 non-redundant bin sets. To dereplicate near-identical genomic bins, Galah (version 0.1.0) was employed to cluster genomes with ≥ 99% average nucleotide identity (ANI) across the four sets of bins generated from each sample type (three individual assemblies and one co-assembly) and select the best genome within each cluster based on completeness and contamination estimates obtained with CheckM [99] (version 1.1.3). Finally, MAGpurify2 was executed to remove putative contaminant contigs within each bin and the final set of bins (referred to as MAGs hereafter) was attained by discarding the bins that did not exhibit the minimum requirements to be classified as medium or high-quality MAGs according to the MIMAG standard [100]. We estimated MAG abundance using CoverM (version 0.3.2, available at https://github.com/wwood/CoverM) in genome mode (parameters: ‘--min-read-percent-identity 0.95 --methods mean’).

### Taxonomic assignment, species clustering and phylogenetic novelty of MAGs

MAGs were assigned taxonomic classifications using GTDB-Tk [101] (version 1.1.0), which uses 120 bacterial and 122 archaeal markers to place the genomes into reference bacterial or archaeal phylogenies (GTDB release 89), depending on which market set was predominant in the genome. In addition to the GTDB-based taxonomy, MAGs were also grouped into *de novo* taxonomy-independent species clusters using a greedy algorithm based on pairwise genome comparisons. Briefly, species clusters were iteratively constructed by grouping pairs of MAGs that aligned across at least 65% of the shorter genome length with at least 95% ANI [16,102,103], according to FastANI [16] (version 1.32) estimates. For each cluster, the highest scoring MAG (where the score was computed as ‘completeness – 5 × contamination’) was chosen as an operational species representative.

The amount of phylogenetic novelty brought by the *campos rupestres* MAGs to different bacterial and archaeal taxa at varying taxonomic ranks (phylum, class, order, family, and genus) was quantified using the phylogenetic gain metric [104], which stands for the added branch length from a subset of genomes within a phylogenetic clade. To measure this metric, the multiple sequence alignments of bacterial and archaeal markers generated by GTDB-Tk were used to reconstruct maximum-likelihood phylogenetic trees with IQ-TREE (version 2.0.3), using the WAG+G substitution matrix and the fast-mode parameter (‘--fast’). The lengths of all branches and MAG-exclusive branches within each evaluated taxon were then measured with the DendroPy [105] library (version 4.4.0).

### Functional annotation and protein clustering

Functional annotation was achieved through the assignment of KO terms[106], Pfam families[107], and TIGRFAM families[108] to genes obtained from the metagenomic assemblies, PLASS’ protein-level assemblies, and MAGs. The genes derived from each of these three sources were annotated separately. Metagenomic assemblies were annotated using the DOE-JGI IMG Annotation Pipeline [109, 110] (v.5.0.15). Protein-level assemblies obtained with PLASS were annotated using KofamScan [111] (KOfam release 2020-12-08, parameters: ‘--e-value 0.01’), a modified version of PfamScan (available at https://github.com/apcamargo/hpc_pfam_search) to assign Pfam (release 33.1) families, and hmmsearch to assign TIGRFAM (release 15.0) families (parameters: ‘--cut_nc’). MAGs were annotated using a modified version of EnrichM (version 0.5.0, available at https://github.com/apcamargo/enrichM), manually modified to update the Pfam (release 33.1) and KEGG (release 2020-05-10) databases and to use pre-established thresholds (‘--cut_ga’ for Pfam, ‘--cut_nc’ for TIGRFAM). Additionally, transporters were identified by querying metagenomic protein sequences against reference transporters from the TCDB database [112] (release 2021/04/07) with MMSeqs2 [113] (release 13-45111) (search parameters: ‘-e 1e-5 --min-seq-id 0.3 -c 0.7 -s 7.5’) and extracting the best hit for each query. Carbohydrate active (CAZy) enzymes were annotated using hmmscan to query metagenomic proteins to dbCAN [114] (version 9) profiles and then filtering the results with dbCAN’s hmmscan-parser script (thresholds: alignment coverage ≥ 35%, E-value ≤ 1e-18).

Proteins predicted from the 16 metagenomic and 12 PLASS assemblies were jointly clustered using Linclust [115] (version 12.113e3). We clustered proteins at multiple identity levels (100%, 95%, 90%, and 50%, using the ‘--min-seq-id’ parameter), requiring that the proteins within a cluster had at least 90% of their sequence length covered by the cluster representative (parameters: ‘--cluster-mode 2 --cov-mode 1 -c 0.9 --kmer-per-seq 80’).

### Read-level gene identification

We used GraftM [116] (version 0.13.1) to identify target genes directly from read sequences and assign them to a taxonomy according to their phylogenetic placement. This approach was employed in cases where we expected the genes to be poorly represented in the assemblies: orthologs of the *Saccharomyces cerevisiae* high-affinity phosphate:H^+^ symporter (*PHO84*), and *Bradyrhizobium* orthologs of the *nifH* gene. In the former case, read-level identification was used because eukaryotic contigs are remarkably difficult to assemble in complex metagenomes, while in the latter this approach was necessary to increase the detection sensitivity of genes encoded by endosymbiotic bacteria.

To identify the *PHO84* orthologs in our reads, we retrieved all UniProt (release 2020_06) protein sequences assigned to the TIGR00887 TIGRFAM family and used them, as well as their taxonomies, as inputs to generate a custom GraftM database. During this process, manual intervention was required to root the phylogenetic tree using phytools (version 0.7-70) ‘midpoint.root’ function. For *nifH*, we retrieved all genes annotated with the K02588 KO term from AnnoTree [69] (doi:10.5281/zenodo.3732466), as well as their taxonomic lineages, and used them to build GraftM databases.

### MAG metabolic potential inference

Due to the inherent incompleteness of MAGs, it is challenging to ascertain their metabolic potential, as it can be uncertain whether the absence of a given gene is due to technical (the gene was not assembled or binned) or biological (the gene is not in the source genome) reasons [117]. Thus, to determine the metabolic potential of the retrieved MAGs, we established a series of criteria based on the presence/absence of key genes, the completeness of multiprotein complexes, and the fraction of genes in KEGG modules. Specifically:

**Photosynthesis:** at least one protein of a photosynthetic reaction center (annotations: K02689, K02690, K08928, K08929, K02703, K02706, K08940, PF00223, PF00124).

**CBB cycle:** at least 60% of the metabolic steps in the KEGG module (M00165) and at least one Rubisco subunit (annotations: K01601, K01602).

**Aerobic CO oxidation:** encode the catalytic subunit of the aerobic carbon monoxide dehydrogenase (*coxL*) (annotation: TIGR02416).

**Nitrogen fixation:** at least two subunits of the *nifHDK* complex (annotations: K02588, K02591, K02586).

**Ammonia oxidation:** at least two subunits of the *amoABC*/*pmoABC* complex (annotations: K10944, K10945, K10946).

**Hydroxylamine oxidation to nitrite:** presence of the *hao* and cytochrome c554 (necessary for proper *hao* function [118]) genes (annotations: K10535, PF13435). As Archaea can produce nitrite from ammonia without a *hao* ortholog [50], we considered that all archaeal MAGs capable of ammonia oxidation were also able to oxidize hydroxylamine.

**Hydroxylamine oxidation to nitrous oxide:** encode at least one gene containing the Cytochrome P460 domain (annotation: PF16694).

### Gene abundance estimation

To quantify gene abundance across all samples, we first used Linclust to dereplicate nucleotide sequences of the genes predicted from the 16 assemblies by clustering them at 100% identity (‘--cluster-mode 2 --cov-mode 1 --min-seq-id 1.00 -c 0.9 --kmer-per-seq 80’). Next, Salmon (version 1.4.0) [119] was executed to map the reads from the 12 metagenomic samples to the dereplicated gene set and estimate read counts and effective lengths for each gene representative in each sample (parameters: ‘--meta’). Last, we quantified gene abundance using the RPKG (reads per kilobase per genome equivalent) metric so that the values are proportional to the expected gene copy number per cell and comparable between different libraries [120]. To do that, we first estimated the average genome size of each sample using MicrobeCensus [120] (version 1.1.1) and then determined the number of genome equivalents (GE) as (Sample read count * 300)/Average genome size, where 300 corresponds to the total length of each read pair. Next, RPKG was computed using the following formula: (*Mapped reads* * 1,000)/(*Effective length* * *GE*). The effective lengths of each gene in each sample were obtained from Salmon’s output.

### Functional comparison between metagenomes

We used RPKG values to evaluate differences in the the carbon, phosphorus, and nitrogen turnover potential between the substrate and root-associated communities. To do that, we first summed the RPKGs of genes associated with the same biological process (e.g.: nitrogen fixation) and then evaluated for systematic differences in abundance between the contrasted conditions using LMMs as follows: *log*10(*RPKG* + 1) = *β* × *N* + H + P +E, where *β* is the regression coefficient, *S* is the environment (substrate or root), *H* and *P* are the random effects of the host plant species and the biological process, respectively, and *E* is a vector of errors.

Gene abundance data was also used to functionally contrast the metagenomes of *V. epidendroides* and *B. macrantha* by identifying (1) KEGG modules and pathways that are significantly enriched in the communities associated with one plant relative to other, and (2) modules and pathways that are enriched among the genes that are similarly abundant in the metagenomes of both plants. To do that, we translated the dereplicated gene set using gotranseq (version 0.2.2, available at https://github.com/feliixx/gotranseq) with the translation table 11 and clustered the resulting proteins at 90% sequence identity (Linclust parameters: ‘--cluster-mode 2 --cov-mode 1 --min-seq-id 0.90 -c 0.9 --kmer-per-seq 80’). These protein-level cluster definitions were then used to aggregate Salmon’s gene-level estimates into cluster-level abundances using tximport’s [121] (version 1.18.0) ‘scaledTPM’ approach. These cluster-level abundances were then imported into a DESeq2 [122] (version 1.30.0) object and between-sample normalization was performed by setting the library size factor as the ratio between the number of GEs in the sample and the median of the number of GEs across all samples, so that the libraries had a similar number of GEs after normalization. This decision was made so that subsequent statistical tests would effectively compare the expected number of gene cluster copies per cell, a biologically meaningful quantity [120, 123].

Highly differentially abundant gene clusters (log2 fold changes ≥ 0.5 or ≤ −0.5) were identified using a Wald test with strict filtering criteria (s-value ≤ 0.005; DESeq2 options: ‘altHypothesis = “greaterAbs”, lfcThreshold = 0.5’). Significant enrichment of KEGG modules and pathways (FDR ≤ 0.05, score > 0) in the communities associated with each plant was appraised using the Kolmogorov-Smirnov test — as implemented in the hypeR library — on lists of gene clusters ranked by apeglm-shrunken [124] log2 fold changes. Conversely, the modules and pathways that are significantly enriched (FDR ≤ 0.05) among the gene clusters with small differences in average genomic copy number between the microbiomes of the two conditions were identified by selecting gene clusters with significant low log2 fold changes (Wald test FDR ≤ 0.05; DESeq2 options: ‘altHypothesis = “lessAbs”, lfcThreshold = 4’) and then performing a hypergeometric test using hypeR. For these analyses, we assigned to each gene cluster the KOs of all the genes within it and then designated KEGG modules and pathways according to the KO-to-module and KO-to-pathway relationships retrieved using KEGG’s REST API (release 97, 2021/01). Substrate and root-associated metagenomes were processed together.

### Protein family copy number comparison between MAGs and GTDB genomes

To appraise potential genomic adaptations of bacteria from the *campos rupestres* regarding P nutrition, we employed phylogenetic regressions, as implemented in the phylolm [125] package (version 2.6.2, function) to test whether genomes retrieved from *campos rupestres* communities have significant changes regarding the number of genes involved in P turnover processes. To account for differences in genome size, we decided to use the genic density (number of genes per Mb) as the response variable using the following formula: *N*\*10^6^/*Genome size* = *β* × *CP* + *E*,where *N* represents the number of genes involved in P turnover processes in the genome, *CP* represents a binary predictor variable that indicates whether the genome was retrieved from the *campos rupestres*, and *E* is phylogenetic covariance-aware vector of errors. In all instances we used the ‘phylolm’ function with the ‘model = “lambda”’ parameter to use Pagel’s λ [126] to measure the phylogenetic covariance.

Functional annotations for all GTDB (release 89) bacterial genomes were retrieved from AnnoTree. To avoid biases that could arise due to different annotation methods used for KO term assignment by AnnoTree and our MAG annotation pipeline, we reannotated our MAGs using EnrichM’s UniProt-based KO assignment with the same thresholds used by AnnoTree.

### Biosynthetic gene cluster identification and clustering

To identify biosynthetic gene clusters (BGCs) regions, we used antiSMASH [127] (version 5.1.2), ignoring contigs shorter than 5 kb. We then used BiG-SCAPE [128] (version 1.0.1) with the ‘--cutoffs 0.5 --clan_cutoff 0.5 0.7 --mibig’ parameters to cluster these regions into gene cluster families (GCFs), and then cluster GCFs into gene cluster clans (GCC).

To investigate the structural organization diversity of siderophore BGCs in more detail, we set aside all the metagenomic BGS regions where antiSMASH identified at least one siderophore cluster (characterized by the presence of a IucA_IucC domain) and executed BiG-SCAPE on then with the ‘--cutoffs 1.0’ parameter to obtain an all-versus-all distance matrix which was then used to hierarchically cluster the siderophore BGCs using the UPGMA algorithm, as implemented in the ‘linkage’ function of the SciPy [129] (version 1.5.4) library. To reduce BGC structural redundancy for visual representation (Figure 6), groups of highly similar BGC regions were identified with SciPy’s ‘fcluster’ function, using the inconsistency criterion and a 0.1 threshold, and only the medoid within each group was selected for graphical representation (23 out of a total of 42 BGC regions). CoverM (version 0.3.2) was used to measure the average coverage of contigs containing representative siderophore BGCs (parameters: ‘--min-read-percent-identity 0.95 --methods mean’).

### Phylogenetic analysis of nitrogenase genes

To assess the diversity of the *nifH* gene we fetched all the metagenomic proteins that were annotated with the K02588 KO term and retrieved selected orthologs belonging to *nifH* groups I, II, and III [45] from UniProt (release 2020_06) (Suppl. Figure 7B, labels in grey). Linclust was then used to cluster identical sequences from the combined set of *campos rupestres* and reference *nifH* (parameters: ‘--cluster-mode 2 --cov-mode 1 --min-seq-id 1.0 -c 0.9 --kmer-per-seq 80’). The dereplicated *nifH* sequences were then subjected to a two-step alignment process: the sequences were first aligned with MAFFT (parameters: ‘--auto’) and poorly aligned proteins were identified and removed from the alignment using trimAl [130] (version 1.4.1) with the ‘-resoverlap 0.3 -seqoverlap 90’ parameters. The remaining sequences were then realigned and unreliable regions were trimmed with trimAl (parameters: ‘-automated1’). Finally, we used IQ-TREE (parameters: ‘-B 1000’) to infer the maximum likelihood tree from the resulting alignment.

To evaluate the hypothesis that the *campos rupestres* Isosphaeraceae *nif* complex was acquired by HGT, we reconstructed a nitrogenase phylogenetic tree using sequences from the *niH* and *nifD* subunits. In addition to the orthologs encoded by the Isosphaeraceae MAGs, we retrieved *niH* and *nifD* sequences from all *nif*-encoding Planctomycetota (AnnoTree r89), from Frankia, and from selected Gammaproteobacteria with high sequence identity to the MAGs’ genes (UniProt release 2020_06) (Suppl. Figure 7C). Next, we generated multiple sequence alignments for the *nifH* and *nifD* sequences using MAFFT (‘-- auto’) and trimmed them with trimAl (‘-automated1’). Finally, IQ-TREE (‘-B 1000’) was employed to infer the maximum likelihood *nifHD* tree using the alignments of both subunits in a partitioned model. To further support the *nif* HGT hypothesis, we used clinker [131] (version 0.0.16) to compare a *nif*-containing Isosphaeraceae contig to the *Pseudomonas stutzeri* A1501 (accession: GCA_000013785.1) *nif* cluster.

### Identification of bacteriophages, proviruses, and their associated hosts

To identify contigs representing bacteriophage genomes or provirus insertions in host sequences, we employed VIBRANT [132] (version 1.2.1) to scan all the metagenome assemblies. The resulting sequences were then processed by CheckV [133] (version 0.7.0) to remove low-quality viral fragments and refine the boundaries between host and provirus regions. Auxiliary metabolic genes were identified in the remaining sequences using KofamScan (KOfam release 2020-05-10). To assign taxonomy to provirus hosts we first removed the predicted viral regions from provirus-bearing contigs and then used MAGpurify2 (version 1.0.0, database available at doi: 10.5281/zenodo.3817702) to classify the provirus-free segments.

## Data availability

16S and ITS amplicon sequencing data are available at the NCBI Sequence Read Archive (SRA) under the BioProject PRJNA522264. The SRA and IMG identifiers associated with the metagenomic data are listed in Suppl. Table 1. The metagenome-assembled genomes used in this work were deposited in GenBank and their accessions are listed in Suppl. Table 2. Supplementary files containing additional processed data are available at https://doi.org/10.17605/OSF.IO/XKDTV.

## Funding

This work was supported by grants from Fundação de Amparo à Pesquisa do Estado de São Paulo (FAPESP) (2016/23218-0) and Coordenação de Aperfeiçoamento de Pessoal de Nível Superior (CAPES) (88881.068071/2014-01). A.P.C. received a scholarship (2018/04240-0) from FAPESP. R.S.C.S received a postdoctoral fellowship (2018/19100-9) from FAPESP. I.R.G. and P.A. are Conselho Nacional de Desenvolvimento Científico e Tecnológico (CNPq) research fellows. The work conducted by the US Department of Energy Joint Genome Institute is supported by the Office of Science of the US Department of Energy under contract no. DE-AC02-05CH11231.

## Notes

### Competing Interest Statement

The authors have declared no competing interest.

