## Supplementary material for "Plant-associated microbiomes promote nutrient turnover in impoverished substrates of a biodiversity hotspot": Suppl.: Supplementary_Note_1.pdf

### Supplementary Note 1: Estimating the sequenced diversity recovery rates in the metagenomic assemblies and MAGs

Due to their remarkable microbial diversity, soil and root associated sequencing data were particularly difficult to assemble into metagenomes represent a significant portion of the total sequence diversity. Therefore, we estimated the recovery rate of our single-sample assemblies by mapping raw reads to the metagenomes and MAGs and we found that metagenomes recovered 16.3% to 42.4% of the total sequence diversity across different samples, whereas the MAGs recovered 10.8% to 39.1%. Metagenome co-assemblies, which were assembled from the combined read set of all samples within each environment, had a minimum recovery rate of 28.2% (Suppl. Table 3).

Assessment of the recovered diversity using sequence matching of 14 different ribosomal protein (RP) markers provided largely concordant results. In addition, clustering RPs with at least 89% identity revealed that at an approximate genus-level divergence the metagenomes and MAGs recovered a much larger fraction of the sequence diversity (Suppl. Table 3), indicating that most of the unassembled sequences belonged to taxa close to those represented by the assembled metagenomes.

To evaluate whether read complexity could have caused the variation in sequence recovery, we estimated the read k-mer diversity of all samples (Suppl. Table 3) and found high and statistically significant negative correlations between the sample diversity and the metagenomic sequence retrieval (mapping rate Spearman's $\rho \approx -0.80$, $p$-value $< 0.01$; RPs Spearman's $\rho \approx -0.79$, $p$-value $< 0.01$), indicating that elevated sequence diversity was the main factor that hindered metagenome assembly.

#### Methods

To quantify how much of each community is represented in the metagenomic assemblies and MAGs two approaches were employed: the rate of read mapping was measured and the presence of single copy markers was determined. The mapping rate of each environment was obtained using Bowtie 2 to map reads from all samples associated with that environment to a concatenation of the four assemblies derived from them and to a concatenation of all MAGs; the fraction of reads that mapped to those references with at least 95% identity (‘--min-read-percent-identity 0.95 --methods relative_abundance’) was then obtained with CoverM in genome mode. For the single copy markers, the recruitment of 14 ribosomal markers in each environment was assessed using SingleM (version 0.13.2, available at https://github.com/wwood/singlem), which identifies these genes in the reads, metagenomic assemblies, and MAGs, and then matches their sequences to obtain the fraction of markers found in the reads that were recovered in the assemblies and MAGs. Genus-level recovery was appraised by allowing SingleM to perform imperfect matches by clustering markers at 89% amino acid identity, which approximates to the genus-level diversity for these markers. Read sequence diversity was computed from quality-trimmed reads with Nonpareil [1] (version 3.3.4) using the k-mer algorithm. Differences in sequence diversity between the metagenomes of the two plants were determined using a linear model, excluding soil and rock samples.
