## Supplementary material for "Plant-associated microbiomes promote nutrient turnover in impoverished substrates of a biodiversity hotspot": Suppl.: Supplementary_Note_2.pdf

### Supplementary Note 2: Annotation-independent assessment of functional novelty through comparative genomics

We quantified functional novelty using a taxonomy-informed approach by comparing 398 of the retrieved MAGs to genomes related at the family level and identifying groups of orthologous genes (orthogroups) that were exclusive to the *campos rupestres* genomes at the family level. Among the 51 families that were analyzed, we found 563 exclusive orthogroups (out of 237,989) in 44 families (median across the 47 families: 6.5), comprising 310 genomes with at least one exclusive orthogroup (median across the 310 genomes: 4), and a total of 4,829 genes (median across the 310 genomes: 8), 4,110 of which were complete. To understand the evolutionary history of these orthogroups, we investigated their distributions across the entire bacterial phylogeny and classified them into four categories according to their likely origin within the family (Suppl. Figure 2A): (1) origination, where orthologs could not be found in any other genome (148 orthogroups); (2) putative horizontal gene transfer (HGT), when orthologs were found in lineages that were inconsistent with the ones in our data (199 orthogroups); (3) lost in other members of the family, when orthologs were found in other families within the same order (18); and (4) other, when the orthogroup was assigned to high taxonomic ranks that were consistent with the family lineage, so it was not possible to determine the precise evolutionary history of the orthogroup (198 orthogroups) (Suppl. Figure 2B). We note that because most of the bacterial diversity has not been sequenced, the number of orthogroups in the origination category is probably overestimated to an unknown degree.

The 148 orthogroups that were inferred to have originated within clades of *campos rupestres* genomes were distributed across 31 bacterial families (median across the 31 families: 2) and were characterized by having an exceptionally low proportion of annotated genes (Suppl. Figure 2C, Suppl. Table 7). Only four of the orthogroups from the origination class contained known structural domains in at least one third of their genes: a Pseudonocardiaceae orthogroup (4 tandemly arrayed genes in the same genome) containing a GntR DNA-binding helix-turn-helix domain (PF00392), which has been associated with plasmid function [1] and was located right next to another plasmid-associated conjugative protein (PF10412); a Streptosporangiaceae orthogroup (7 genes distributed across 4 genomes) that possessed a STAS domain, which is involved in the control of the activity of antisigma-factors; a Xanthobacteraceae orthogroup (4 genes in the same genome, 3 of which were located near each other) with a PEP-CTERM domain (PF07589); and an Acetobacteraceae orthogroup (3 genes in the same genome) with a PepSY domain (PF13670). Both the Xanthobacteraceae and Acetobacteraceae orthogroups were comprised of proteins containing the Sec/SPI signal peptide, which was consistent with other proteins harboring PEP-CTERM and PepSY domains. In addition to the protein domains, we also found that 38 orthogroups from the origination class (≈ 25.68%) had at least one third of their genes containing a signal peptide, 31 (≈ 20.95%) had at least one third of their genes containing at least one transmembrane helix domain, and 5 (≈ 3.38%) had at least one third of their genes within biosynthetic gene clusters (BGCs). Interestingly, there seemed to be an enrichment in secreted peptides among the origination orthogroups, as the proportion of orthogroups annotated with signal peptides across all inferred orthogroups (≈ 17.13%) was 1.5× lower than in the origination class.

Because functionally novel genes tend to be lineage-specific [2], we also investigated the genes without identified functional domains within orthogroups from the origination class. We found that 145 of these orthogroups (97.97%) had at least one third of their genes without any assigned KO, TIGRFAM, or Pfam, which was approximately 3× more than the proportion across all the identified orthogroups (≈ 32.74%). Furthermore, we used sensitive HMM-HMM comparisons to search for remote homologs — that, due to low sequence identity, could be detected by standard protein search methods — of the origination orthogroups and found that only 35 (≈ 23.65%) of them had significant hits to protein clusters of the UniClust30 HMM database [3] (all belonging to bacteria, and 31 containing a single sequence), and that 30 matched profiles without known function. Aside from four protein domains mentioned above, this search found one distantly related hydrogenase maturation protein. We hypothesize two scenarios for some of our orthogroups from the origination class having distant orthologues in other bacteria: they could have been horizontally transferred from a microorganism that is still absent in protein databases, or they could have substantially diverged from the closest known orthologs due to strong selection.

The 199 orthogroups that were assigned to the putative HGT class were distributed across 36 families (median across the 36 families: 36) and were also enriched in protein containing signal peptides (≈ 23.12%) and proteins without functional domains (≈ 60.80%) (Suppl. Figure 2C, Suppl. Table 7). Analysis of the orthogroups containing proteins with known function revealed that the putative HGT orthogroups contained several genes associated with mobile genetic elements (MGEs), such as transposases, integrases/recombinases, membrane fusion proteins, and group II intron reverse transcriptases. Furthermore, these orthogroups were also enriched in genes that are commonly found within MGEs and can confer a selective advantage to the bacterial population, such as antibiotic synthesis genes (clorobiocin, avermectin, and enediyne) and xenobiotic degradation genes.

Altogether, these results showed that the *campos rupestres* MAGs featured a high degree of functional novelty. We expect that some of the genes of unknown function — mainly the ones within BGCs, that produce secondary metabolites, and the ones containing signal peptides — could be involved in novel forms of plant-microbiome interactions.

### Methods

Proteins from the *campos rupestres* MAGs were compared to those of related organisms to infer the evolutionary relationships between their genes and define clusters of proteins that share a common ancestor (orthogroups). To do that, protein sequences from species that belonged to families to which at least one *campos rupestres* MAG was assigned were retrieved from GTDB (release 89); only families encompassing at least three genomes — other than the MAGs generated in this study — were investigated. OrthoFinder [4] (version 2.3.11) was then employed to perform all-versus-all pairwise comparisons between each family's proteins, cluster them into orthogroups, and generate count matrices holding the number of genes within each ortogroup per genome. Orthogroups inherited functional annotations (hypothetical protein, KO, TIGRFAM, Pfam, signal peptide, transmembrane helix, and BGC) from their genes if the annotation was shared by at least one third of them.

To identify orthogroups that were exclusive to the *campos rupestres* MAGs, tspex (version 0.6.2, available at https://github.com/apcamargo/tspex) was used to calculate the specificity measure (SPM) of each orthogroup from the matrices of orthogroup counts. Additionally, to reduce the false positive rate, the number of genes within each family was accounted for by modeling the count data using binomial distributions. Only orthogroups with a minimum of three genes, SPM = 1.0 and $p$-value < 0.001 were considered enriched in the MAGs under investigation. The $p$-value threshold was established by iteratively reducing its value until there was no significant correlation between the number of GTDB genomes in the family and the number of exclusive orthogroups identified (Spearman's $\rho = 0.055$, $p$-value $\approx 0.7$).

The identified exclusive orthogroups were then classified taxonomically using MMSeqs2 [5] (version 12.113e3) and taxopy (version 0.5.0, available at https://github.com/apcamargo/taxopy) softwares. First, each protein was assigned to a taxon in the MAGpurify2 taxonomic database (doi: 10.5281/zenodo.3817702, based on

GTDB release 89) using MMSeqs2's 2bLCA algorithm ('-e 1e-05 --cov-mode 0 -c 0.5 --lca-mode 3'), and then, to assign taxa to orthogroups, the taxonomies of their proteins were aggregated using the majority vote approach, as implemented in taxopy. If no taxonomy was assigned to any of the proteins within the orthogroup (i.e.: they had no significant matches with the database), the process was repeated using the UniRef100 database [6] (release 2020_06). Finally, the orthogroups were classified into four categories according to their likely origin within the *campos rupestres* families: (1) if no protein within the orthogroup had a hit in the MAGpurify2 or UniRef100 databases, the orthogroup was classified as "origination"; (2) if the orthogroup was assigned to a lineage that was discordant with the *campos rupestres* family lineage at any rank, the orthogroup was classified as "putative HGT"; (3) if the orthogroup was assigned to the order to which the *campos rupestres* family belongs, it was classified as "lost in other genera of the family" class; (4) if the orthogroup was assigned to a high-rank taxon (class, phylum, domain, or root) that was not discordant with the *campos rupestres* family lineage, it was classified as "other", as it was not possible to infer its probable evolutionary history. These classifications assumed a single horizontal gene transfer or loss event.

We conducted HMM-HMM searches to find remote orthologs of the orthogroups within the "origination" class using the HHblits [7] (version 3.3.0) software. First, the complete proteins within each orthogroup were aligned using MAFFT [8] (version 7.464) with the '--auto' parameter and then the multiple sequence alignments were converted into the A3M format using the 'reformat.pl' script of the HH-suite. Finally, each orthogroup A3M alignment was subjected to an iterative HMM-HMM search against the UniClust30 HMM database[3] (release June 30, 2020) and we accepted the first hits with an estimated homology probability ≥ 90%, E-value ≤ $10^{-5}$, and target coverage ≥ 50%.
