## Supplementary material for "Plant-associated microbiomes promote nutrient turnover in impoverished substrates of a biodiversity hotspot": Suppl.: Supplementary_Note_3.pdf

### Supplementary Note 3: Assessing populational divergence of bacterial species shared by the *V. epidendroides* and *B. macrantha* microbiomes

We also assessed microbiome differentiation by evaluating intra-species populational divergence. To do that, we investigated single nucleotide variant (SNV) sites of populations of species that were highly abundant in the microbiomes of both plants, evaluating differences in allele frequencies, genic nucleotide diversity, and linkage disequilibrium.

In total, we analyzed seven genomes (from 3 Eremiobacterota and 4 Proteobacteria) and found that, even though the *V. epidendroides* and *B. macrantha*-associated populations mostly exhibited differentiation regarding the criteria listed above (Suppl. Figure 4A–C), there were no systematic patterns that distinguished the populations associated with the two plants. Instead, we found that divergences between the pairs of populations seem to be tied to the bacterial taxonomy, rather than the host plant or substrate.

Lastly, we inquired whether the observed differentiation could be attributed to processes other than neutral evolution, such as natural selection or changes in population size. Therefore, we measured the Tajima's D statistic for all genes in each population and noticed that for many populations the distribution was shifted from 0 (Suppl. Figure 4D), indicating that some form of non-neutral evolutionary process might have driven the intra-species divergences between the pairs of populations. Interestingly, some species showed diverging deviations from zero in different populations, suggesting that distinct evolutionary processes led to the differentiation between the populations associated with each plant.

### Methods

To appraise intra-species sequence diversity, recombination dynamics, and measure the genetic divergence of same-species populations associated with the two plant species under investigation, single nucleotide variant (SNV) profiles were obtained for MAG species clusters that were high-abundant in samples associated with both *V. epidendroides* and *B. macrantha*. To carry this out, we built two read sets by concatenating all six FASTQ files from samples associated with each plant (that is, all the substrate and external root sequencing samples associated with the same plant) and then we mapped these sets to each assembly using Bowtie 2. The resulting BAM files were then used to measure the joint coverage of species-representative MAGs (sum of the coverages of all MAGs within the species cluster) in each aggregated sample (using CoverM with the '--min-read-percent-identity 0.95' parameter). We then selected seven species-representative MAGs for further investigation as they exhibited average coverage ≥ 10 in the aggregated read sets of both plants.

Next, inStrain [1] (modified version, available at https://github.com/apcamargo/inStrain) was employed to analyze the read mapping profiles of these seven genomes (parameters: '--filter_cutoff 0.92') by calling SNVs, based on the observed polymorphism frequencies among mapped reads, and measuring linkage disequilibrium (quantified by the $r^2$ metric) using information from read pairs that linked two SNV loci. To compare populations of the same species associated with the two different plants the following properties were assessed: (1) the genetic differentiation with respect to differences in allele frequencies, (2) population-level nucleotide diversity, (3) linkage disequilibrium decay, and (4) deviations from the neutral evolution model. Genomic divergence between populations was quantified from genic allele frequencies in SNV sites using the Hudson estimator [2] of the fixation index (Fst), as implemented in the 'hudson_fst' function of the scikit-allel library (version 1.3.1). Gene-level and genome-level estimates of Fst were attained using the ratio of averages approach [3]. Gene-level estimates of nucleotide diversity (π) for each population were obtained by calculating π for all sites above the coverage threshold (using the 'mean_pairwise_difference' function of scikit-allel) and averaging the values over genes.

Linkage disequilibrium decay profiles were computed across 20 bp windows, averaging $r^2$ values calculated from pairs of sites whose distances were within the same window. Finally, gene-level Tajima's D estimates for each population were calculated from allele frequencies from all sites above the coverage threshold (using the 'tajima_d' function of scikit-allel).
