## Supplementary material for "Plant-associated microbiomes promote nutrient turnover in impoverished substrates of a biodiversity hotspot": Suppl.: Supplementary_Note_4.pdf

### Supplementary Note 4: The carbon turnover potential of *V. epidendroides* and *B. macrantha* microbiomes

Bacteria can obtain carbon from plants indirectly by breaking down the carbohydrates in organic matter with carbohydrate active enzymes (CAZy), such as glycoside hydrolases, polysaccharide lyases, and carbohydrate esterases. We found that CAZy were encoded by multiple taxa (Suppl. Figure 5B) and that they were also enriched in the root-associated microbiomes when compared to the soil and rock communities (Suppl. Figure 5D, LMM *p*-value $< 0.005$; $\omega^2 \approx 0.17$). Surprisingly, we found that CAZy were significantly more abundant in the *B. macrantha* microbiome and the rocks than in the *V. epidendroides* rhizosphere and the soil (LMM *p*-value $< 0.005$; $\omega^2 \approx 0.19$), despite the scarcity of organic matter in the rocks [1].

To better understand the mechanisms of carbon acquisition used by microorganisms that grow over the nutrient-impoverished rocks, we examined the abundances of key enzymes involved in multiple pathways of autotrophic carbon assimilation and found that the Rubisco gene (*rbcL*), which is a hallmark of the Calvin–Benson–Bassham (CBB) cycle, was present in all evaluated metagenomes and was encoded by several phyla (Suppl. Figure 5C). Evaluation of Rubisco abundance revealed that there was no significant difference between the *V. epidendroides* and *B. macrantha* microbiomes (Suppl. Figure 5E) — which was supported by the fact that the CBB cycle pathway was enriched among the genes with similar average genomic copy numbers between in microbiomes of the two plants (FDR $<$ 0.001) — and that 8 out of the 10 families with the highest average *rbcL* abundances were significantly enriched among the set of ASVs that were shared between these plants (Suppl. Table 10), indicating that autotrophy was similarly pervasive in the soil and rock communities despite the difference in organic matter availability.

Because the CBB cycle consumes energy to fix carbon dioxide into organic molecules, we further evaluated the autotrophic features of those communities by estimating the abundances of genes involved in energy generation, either via photosynthesis (PSI and PSII genes) or via carbon monoxide oxidation (*coxL*). We determined that even though both traits were found in the microbiomes of *V. epidendroides* and *B. macrantha*, the communities had different approaches for energy generation as the genes for both photosystems were much more abundant in the *B. macrantha* and rock microbiomes (Suppl. Figure 5E, PSI LMM $p$-value $< 0.05$; PSI $\omega^2 \approx 0.44$; PSII LMM $p$-value $< 0.005$; PSII $\omega^2 \approx 0.61$). This contrast was further backed by significant enrichment in photosynthesis-related processes — such as the biosynthesis of the photosystem types I and II, and antenna proteins (FDR < 0.01) — in these communities. To appraise the autotrophic potential of the studied microbiomes at the genome-level we identified a total of 9 photosynthetic and 32 carbon monoxide-oxidizing MAGs (3 genomes encoded both functions) from diverse taxa, including novel families (Suppl. Figure 5F, Suppl. Table 11).
