## Supplementary figures and images for "Plant-associated microbiomes promote nutrient turnover in impoverished substrates of a biodiversity hotspot"

### Supplementary_Figure_1.pdf

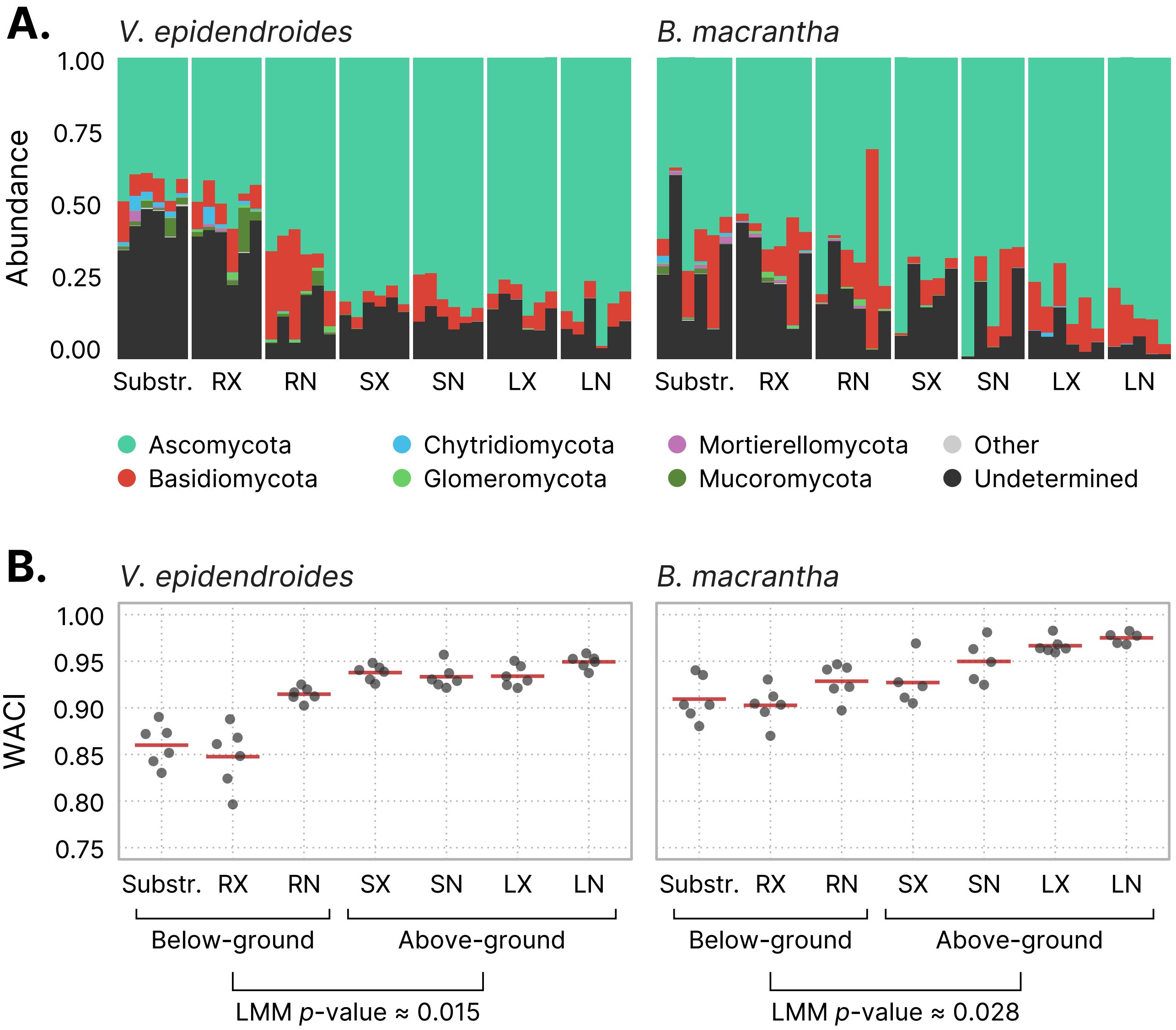

### Supplementary_Figure_2.pdf

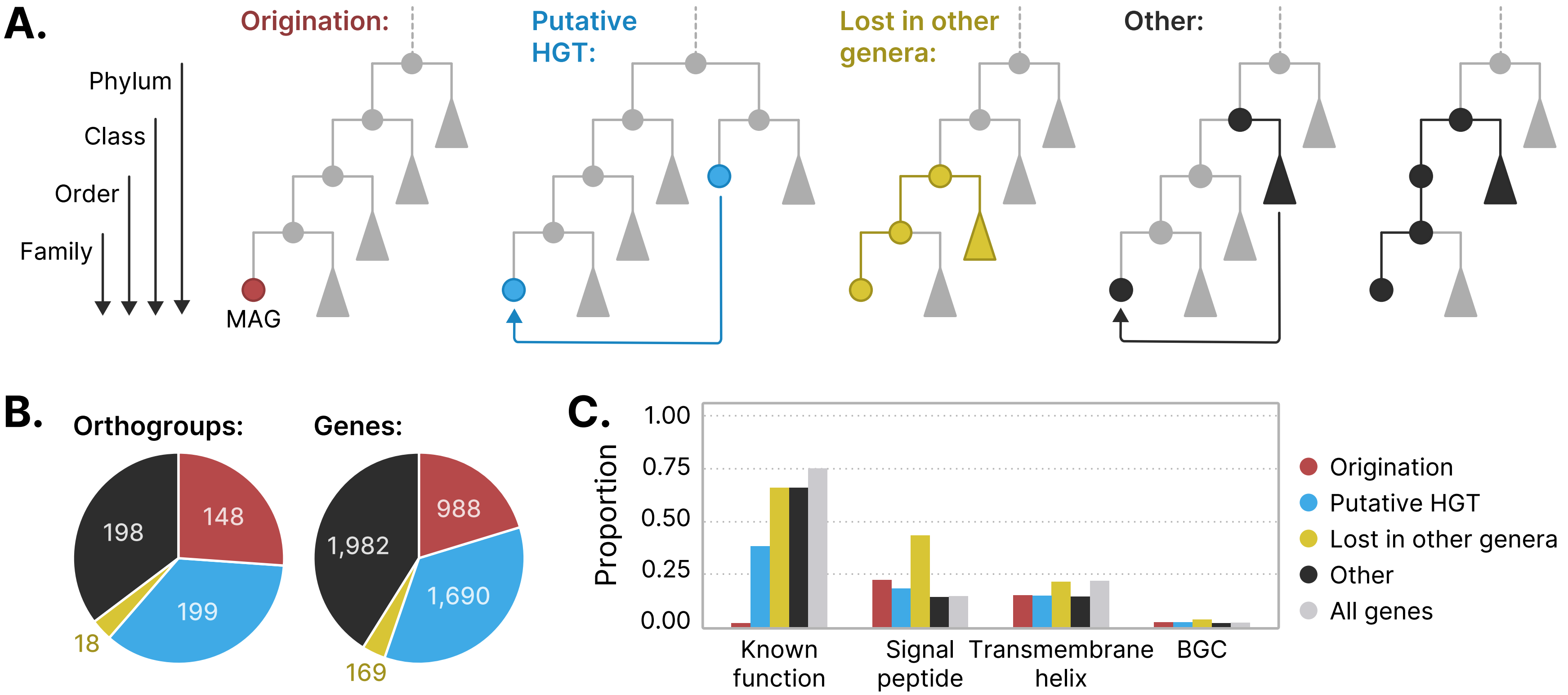

### Supplementary_Figure_3.pdf

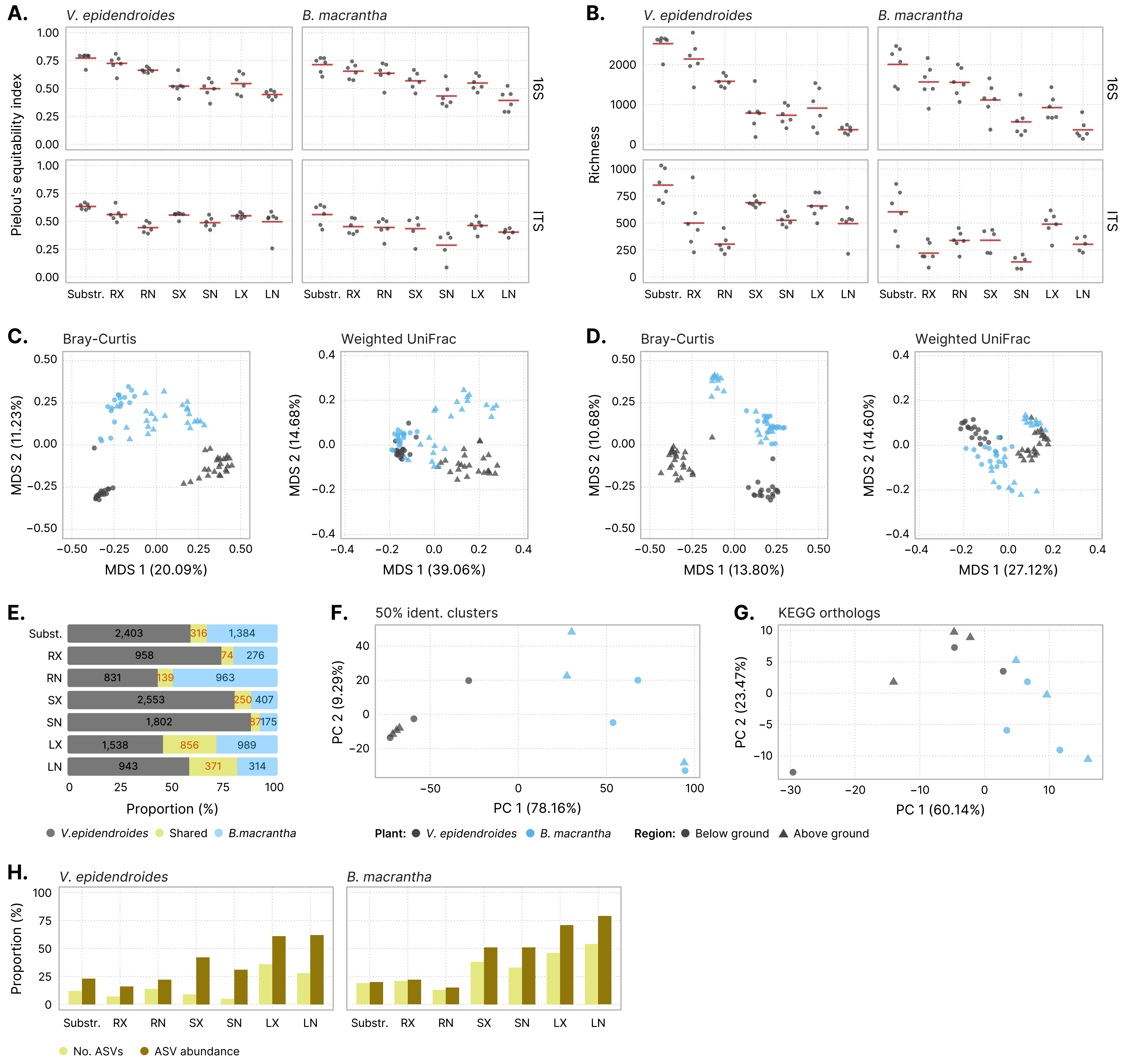

### Supplementary_Figure_4.pdf

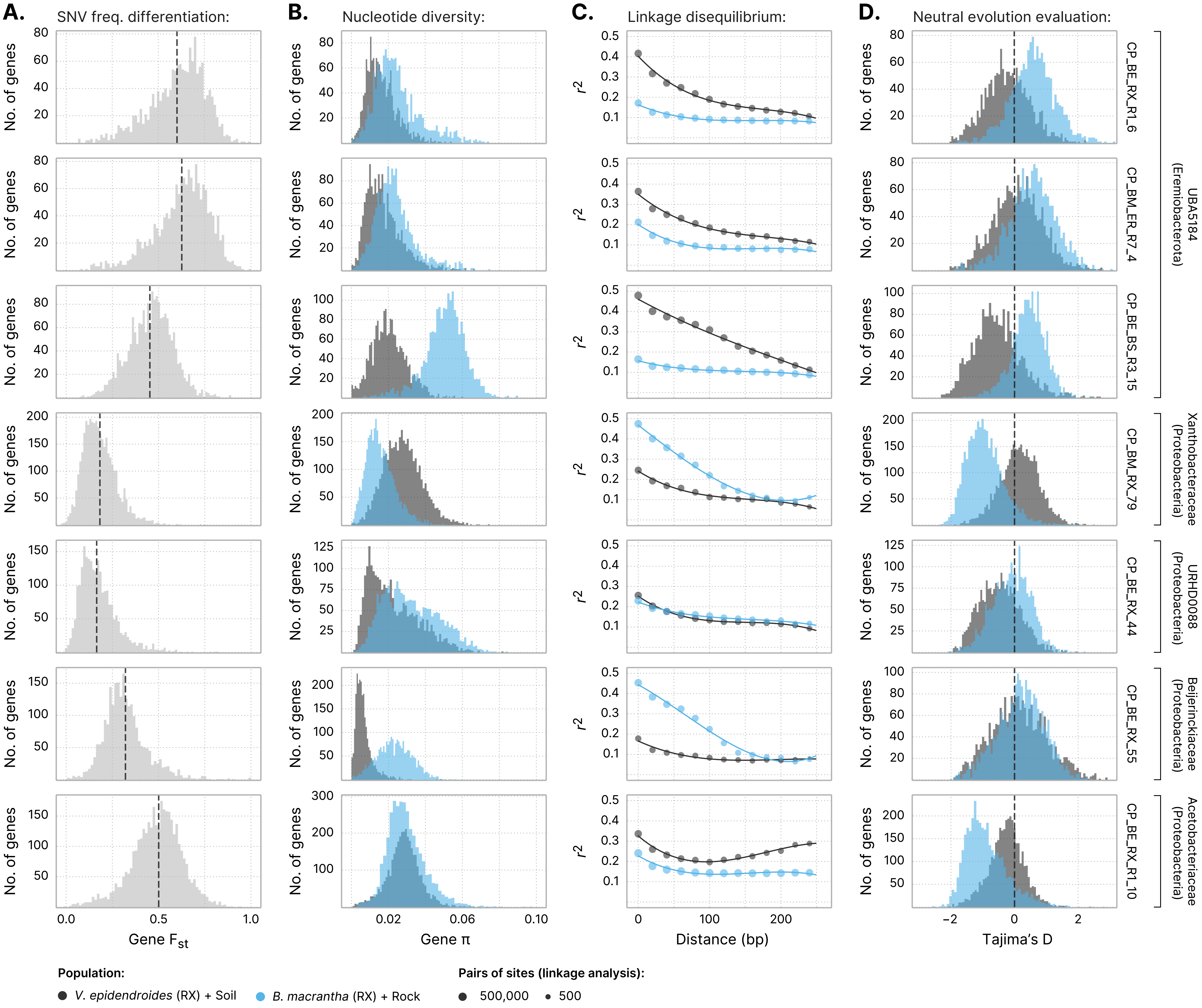

### Supplementary_Figure_5.pdf

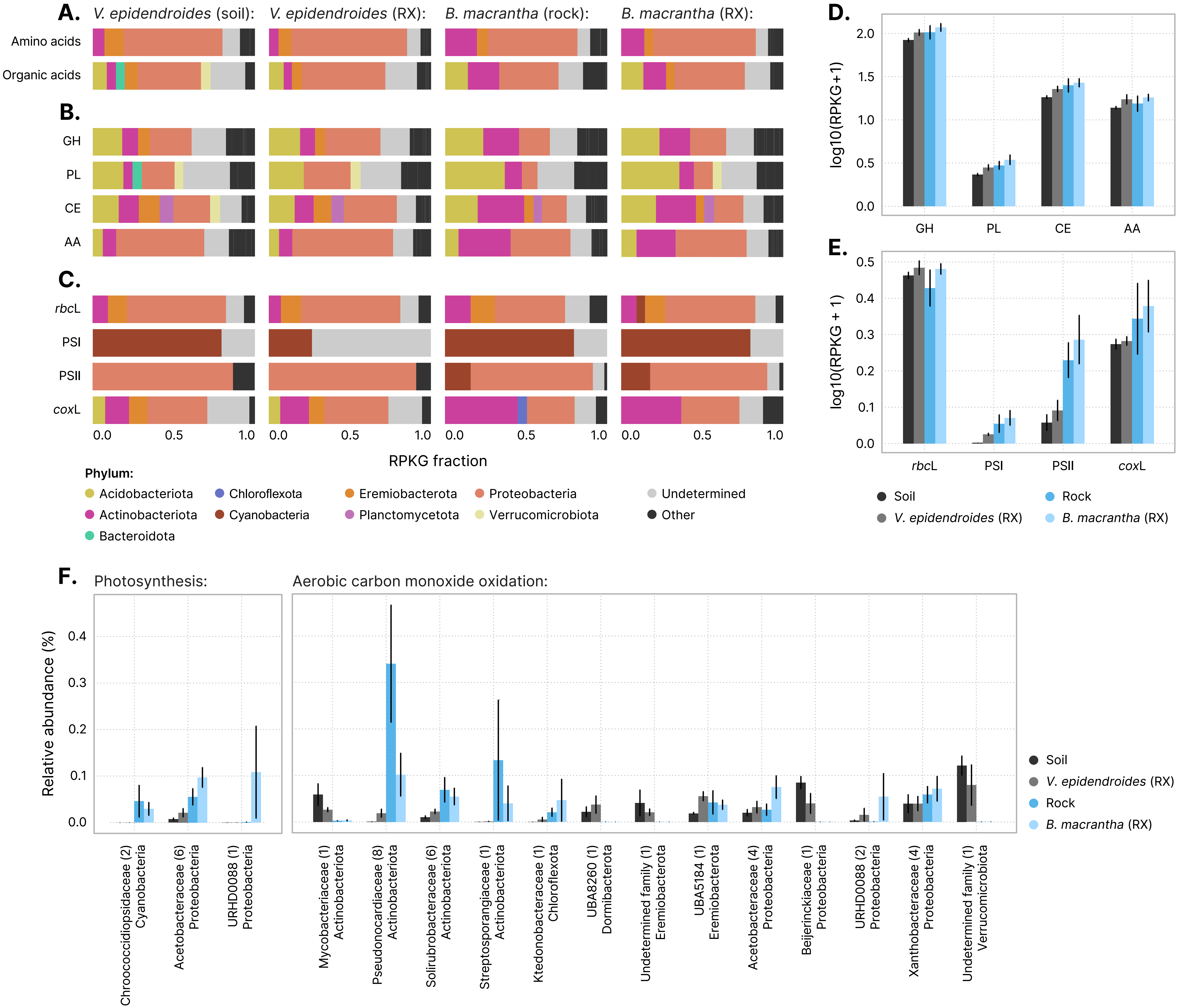

### Supplementary_Figure_6.pdf

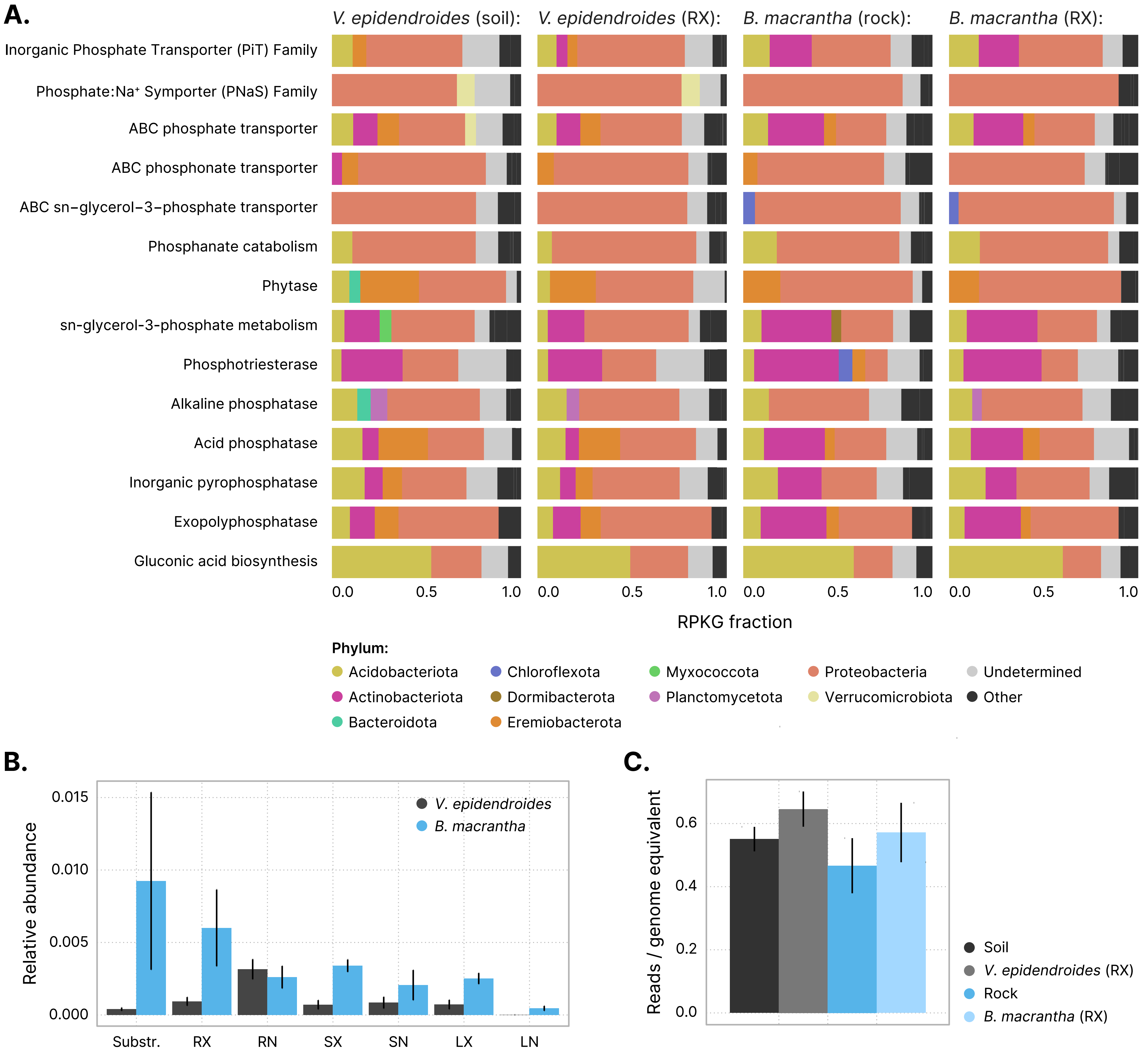

### Supplementary_Figure_7.pdf

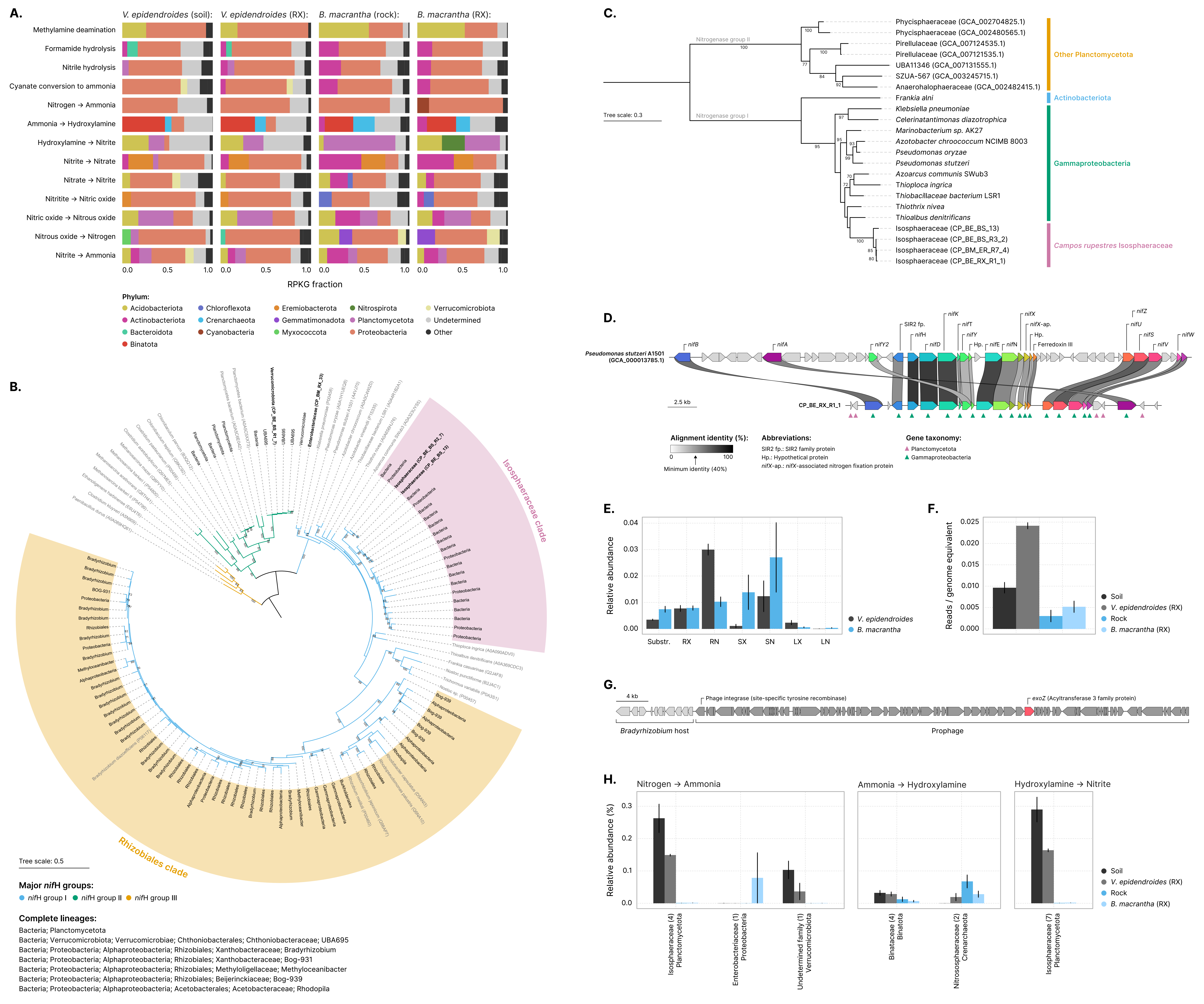
